# CoCoBin: Graph-Based Metagenomic Binning via Composition–Coverage Separation

**DOI:** 10.1101/2025.08.27.672549

**Authors:** Khuanwara Potiwara, Duangdao Wichadakul

## Abstract

**Motivation:** Metagenomic binning is a critical step in metagenomic analysis, aiming to cluster contigs from the same genome into coherent groups. In contemporary workflows, most binning tools begin with the assembly of shotgun metagenomic sequencing data. The assembled contigs are then grouped into bins representing individual microbial genomes or species, typically using taxonomy-independent methods. Although several methods exist, metagenomic binning remains a challenging yet mandatory task, particularly in the context of complex and highly diverse microbial communities.

**Results:** We propose CoCoBin, a novel metagenomic binning tool explicitly designed for the effective binning of metagenomic contigs. In this study, we introduced an innovative approach for calculating contig similarity by separating composition and coverage information. The method begins by (1) assigning contigs into a cluster based on length ranges, (2) calculating contig similarity based on composition features (e.g., k-mer frequencies), and (3) calculating contig difference based on coverage features. These similarity measures are then integrated to construct a graph, where nodes represent contigs and edges represent the similarities between them. Finally, the Louvain algorithm is applied to the graph to cluster closely related contigs. CoCoBin was compared against several state-of-the-art binning tools: BusyBee Web, CONCOCT, MaxBin 2.0, MetaBAT 2, and MetaDecoder on nine simulated datasets, five mock community datasets, and one real dataset. The AMBER tool used to evaluate the binning results across all datasets shows that CoCoBin achieved the best performance regarding the number of bins identified, followed by its performance on the F1 score.

**Availability:** The source code of CoCoBin is available at https://github.com/cucpbioinfo/CoCoBin

**Supplementary information:** Supplementary data are available at Bioinformatics online.

## Introduction

Bioinformatics analysis in microbiome research (Hao et al. 2017) is advancing rapidly, with ongoing development and enhancement of computational tools to characterize the composition and taxonomy of microbial communities. The microbiome (Amon and Sanderson 2017) refers to the community of microorganisms present in a specific environment, including bacteria, archaea, fungi, algae, and small protists. In human contexts (Gilbert et al. 2018), it refers to the microorganisms that inhabit particular regions of the body, such as the skin or gastrointestinal tract. These microbial communities are dynamic and undergo alterations influenced by various environmental factors such as physical activity, dietary choices, medications, and other exposures. Although microbes are too small to be visible, they play a significant role in promoting human health and well-being. They defend us from harmful pathogens, support the development of the immune system, and facilitate the digestion of food to generate energy (Lloyd-Price et al. 2016).

Metagenomics (Sleator et al. 2008) studies the structure and function of microbial genomes within samples from human clinical, animal, and environmental sources to detect and discover variations of microbes under specific conditions. Metagenomic binning (Han et al. 2025) aims to identify known and unknown microorganisms within an environmental sample based on metagenomic sequences. Metagenomic binning (Herazo-Álvarez et al. 2025; Yue et al. 2020) can be divided into two categories: (1) taxonomy-dependent and (2) taxonomy-independent methods. Taxonomy-dependent methods (Madival et al. 2022) rely on the similarity between reads and sequences in reference databases, which are often limited in number. In contrast, taxonomy-independent binning (Sinha et al. 2022) infers groups or bins of reads in a dataset solely based on their similarity, without comparing them to a database. Taxonomy - independent methods employ sequence composition data, machine learning, and statistical approaches to perform binning without relying on reference genomes. These methods are further categorized into composition (k-mer frequency) based, abundance (contig coverage) based, and hybrid methods (combining both the k-mer frequency and coverage features) (Sedlar et al. 2017). For example, state-of-the-art metagenomic binning tools, such as BusyBee Web (Schmartz et al. 2022), CONCOCT (Alneberg et al. 2014), MaxBin 2.0 (Wu et al. 2016), MetaBAT 2 (Kang et al. 2019), and MetaDecoder (Liu et al. 2022) operate without relying on reference databases. Instead, they utilize composition or coverage features of sequences, or a combination of both, to cluster genomic fragments.

In the current era, most metagenomic binning tools typically begin by assembling shotgun metagenomic data. Following assembly, the next step is to group or bin the sequences into clusters representing individual microbial genomes or species based on taxonomy - independent methods. Shotgun metagenomics (short-read sequencing) (Sharpton 2014) involves sequencing DNA derived from a complex mixture of genomes within a microbial community using next-generation sequencing (NGS) technologies, with read lengths ranging from 100 to 300 base pairs (bps). De novo metagenome assembly (Lipovac and Križanović 2023) is the process of reconstructing the genetic sequences of microorganisms solely from DNA sequencing reads without relying on reference genomes.

BusyBee Web (Schmartz et al. 2022) is a web application. It employs bootstrapped supervised binning (BSB) for metagenomic sequencing datasets. BSB does not rely on reference databases and utilizes both unsupervised and supervised machine learning techniques, focusing on genomic signatures like oligonucleotide frequencies. The unique aspect of BusyBee Web is its combination of unsupervised and supervised methods, which creates training data from the input itself rather than relying on existing references. CONCOCT (Alneberg et al. 2014) directly groups all contigs into genomic bins. It merges a coverage vector and a tetramer frequency vector into a unified vector for each contig, employs principal components analysis (PCA) to reduce dimensionality, and uses the Gaussian Mixture Model (GMM) fit with a variational Bayesian approximation for contig binning. MaxBin 2.0 (Wu et al. 2016) utilizes an Expectation–Maximization algorithm to maximize the probability that a contig belongs to a particular bin based on tetranucleotide frequencies and coverage profiles. MetaBAT 2 (Kang et al. 2019) utilizes a clustering algorithm that operates on a graph structure. It begins by computing pairwise distances between all pairs of contigs using composition and coverage information. It derives the composition feature from an empirical posterior probability determined from a collection of reference genomes. Contigs are then assigned into bins based on their distances, with similarity scores determining the connections between contigs in the graph. The binning process employs a label propagation algorithm for graph partitioning. MetaDecoder (Liu et al. 2022) is constructed as a two-layer model. At the first layer, the k-mer frequencies and coverages of all contigs are merged as inputs to the GPU-based modified Dirichlet process Gaussian mixture model (DPGMM) for preliminary clustering, aiming to dissolve small clusters and reassign contigs to the other clusters to avoid over-segmentation. The second layer consists of a semi-supervised k-mer frequency probabilistic model and a modified Gaussian mixture model for modeling coverage based on single-copy marker genes.

According to the study, current metagenomic binning tools still exhibit errors in predicting the number of groups within metagenomic datasets. To address this limitation, we introduce CoCoBin, a metagenomic binning tool designed explicitly for binning metagenomic contigs. CoCoBin improves the contig clustering accuracy by introducing an innovative approach for calculating contig similarity, which separates composition and coverage information. The similarity is then used to construct a graph, which is subsequently clustered using a community detection methodology to achieve effective binning.

## 2. Methods and Materials

The CoCoBin workflow, as shown in Fig. 1, is mainly divided into the following steps: 1) Assembling reads into contigs; 2) Extracting composition features; 3) Computing contig similarity, which involves grouping contigs based on length ranges and calculating similarity and differences based on composition and coverage features; 4) Structuring complex networks; and 5) Clustering.

**Fig. 1.**
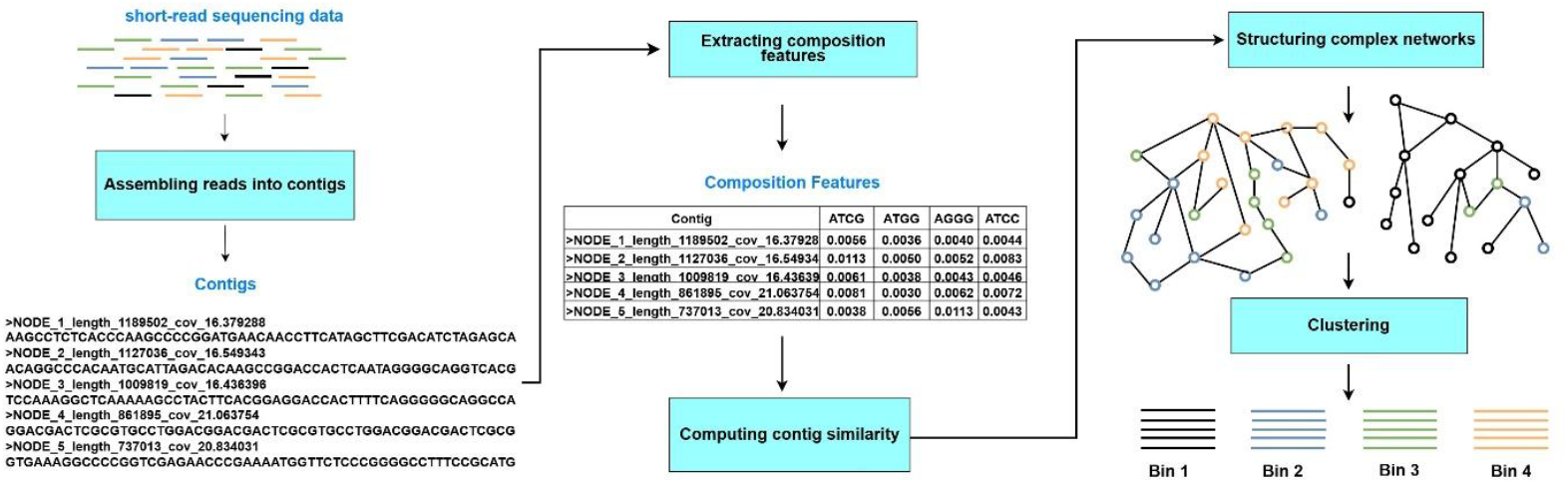
Overview of CoCoBin workflow

### 2.1 Assembling reads into contigs

The short-read sequencing data were *de novo* assembled into contigs using metaSPAdes (Nurk et al. 2017) through Galaxy (Community 2022) with default parameters (selecting the ‘auto’ k-mer detection option).

### 2.2 Extracting composition features

Binning tools such as CONCOCT, MaxBin 2.0, MetaBAT 2, and MetaDecoder require contig lengths greater than 1,000 base pairs (bps) for effective operation. Hence, our CoCoBin employed a default length threshold of 1,000 bps to filter out shorter contigs. Analyzing composition patterns (k-mer occurrences) can categorize the contigs, as different organisms typically exhibit distinct patterns of k-mer occurrences (Alneberg et al. 2014). For the binning process, we relied on the composition information of the contigs, which referred to the frequencies of k-mers (short subsequences of length k). We utilized the iLearn tool (Chen et al. 2019) to generate composition features based on tetranucleotide frequencies (4-mers) of the contigs, resulting in 256 dimensions. To reduce dimensionality, we applied the reversed complement method, treating k-mers and their reverse complements as equivalent, thereby reducing the dimensions from 256 to 136. We then normalized the composition features across contigs to account for the varying lengths of the contigs.

### 2.3 Computing contig similarity

Assessing contig similarity within complex metagenomes presented a significant challenge in microbiome research. Although several binning tools are available to tackle this issue, a gap remains for enhancing accuracy and comprehensiveness. In this study, we introduced a novel methodology for calculating contig similarity that considered separating composition and coverage information and adapted the contig nodes connecting by similarity to the Louvain algorithm (Blondel et al. 2008). The framework for computing contig similarity, as shown in Fig. 2, was divided into steps: 1) Assigning contigs into a cluster based on length ranges, 2) Calculating contig similarity based on composition features, and 3) Calculating contig difference based on coverage feature.

**Fig. 2.**
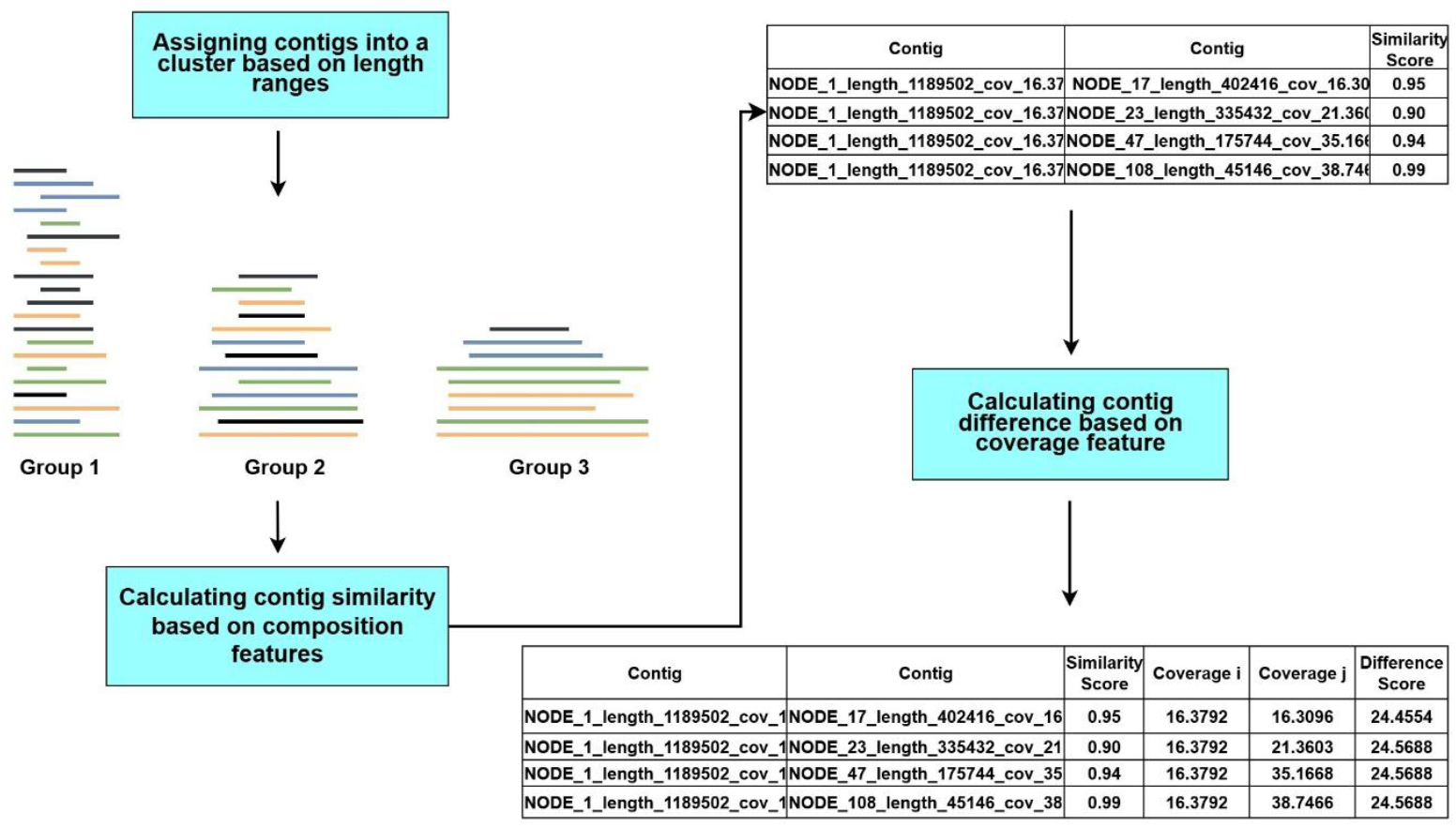
Computing contig similarity workflow

#### 2.3.1 Assigning contigs into a cluster based on length ranges

In this step, the analysis of contig distribution across each dataset revealed a high density of contigs within the length range of 1,000 to 3,500 base pairs (bps), with significantly lower densities for contigs exceeding this length range. The number of contigs and their distribution across organisms for each dataset are presented in Supplementary Tables S2-S15. Based on the distribution of contig length ranges, we proposed assigning each contig to a group according to its length range. This grouping served as a parameter for calculating contig similarity and analyzing microbial communities. The contigs were divided into three groups based on the following length ranges:

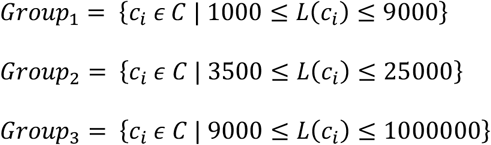

Where *C* = {*c*_1_, *c*_2_, …, *c*_*n*_} is the set of all contigs.

*L*(*c*_*i*_) is the length of contig *c*_*i*_.

#### 2.3.2 Calculating contig similarity based on composition features

CoCoBin, separates the analyses of composition features from coverage feature. In this step, after assigning contigs to clusters based on length ranges (Section 2.3.1), we used only composition features—specifically, k-mer frequency—to calculate contig similarity using the cosine similarity method, as outlined in Equation (1). Each contig was compared only to others within its own cluster. Contig pairs were considered similar if their similarity score was greater than or equal to a predefined threshold (0.85), derived from the experiments. After computing composition-based similarities within each cluster, we further refined the pairwise relationships by incorporating coverage differences (Section 2.3.3).

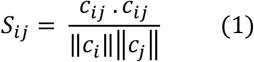

Where *S*_*ij*_ is the similarity between contigs *i* and *j*.

*c*_*i*_, *c*_*j*_ are k-mer composition of contigs *c*_*i*_ and *c*_*j*_ respectively.

#### 2.3.3 Calculating contig difference based on coverage feature

The coverage information of contigs in a metagenomic sample reflected the abundance of microorganisms (Alneberg et al. 2014). It could, therefore, be used to determine the source of each contig (Liu et al. 2022). Contigs from the same source population were expected to have similar coverage, whereas contigs from different source populations often exhibited differing coverage levels (Strous et al. 2012). In this step, we further considered contig pairs passing the similarity score threshold based on composition features from Section 2.3.2 to determine the allowable difference in coverage values between contigs derived from the same genome. Specifically, we investigated the extent of coverage difference that could be tolerated while maintaining accurate binning performance. By analyzing coverage differences between pairs of contigs, we found that a coverage difference of no more than 1.5-fold between contig *i* and contig *j* was optimal for accurately identifying the number of bins (i.e., groups of contigs from the same source population), as outlined in Equation (2). We extracted the coverage feature from the output of the MetaSPAdes tool, and we detailed clustering experimental results in Supplementary Tables S16-S19.

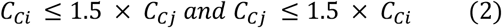

where *C*_*Ci*_ is the coverage information of contig *i*, and

*C*_*Cj*_ is the coverage information of contig *j*.

### 2.4 Structuring complex networks

In this step, we constructed a graph of contig pairs from all groups together with a similarity score greater than or equal to a predefined threshold and a coverage difference of no more than 1.5-fold. The graph was denoted as *G*(*V, E*), where each vertex *ν*_*i*_ represented a distinct contig, and edges *e*_*ij*_ between vertices *ν*_*i*_ and *ν*_*j*_ represented the similarity and difference measures based on composition and coverage features between the contig pairs. We used NetworkX (Hagberg et al. 2008), a Python package for constructing and analyzing complex graphs.

### 2.5 Clustering

We applied the Louvain algorithm based on modularity optimization to identify bins in a contig graph. The algorithm consisted of four steps as follows.

**Step 1:** Initialization – Each contig was initially assigned to its community.

**Step 2:** Modularity Optimization – A common method to detect communities was to find a partition of the vertex set that maximized an optimization function. The quality of the resulting partitions was often measured by the so-called modularity of the partition. The algorithm evaluated the gain in modularity resulting from moving a contig *i* from its current community to the community of contig *j*. Contig *i* was then placed in the community that yielded the highest positive gain in modularity. If no positive gain was possible, contig *i* remained in its original community.

**Step 3:** Community Aggregation – A new network was built where each contig represented a community identified in the step 2.

**Step 4:** Iteration – Steps 2 and 3 were applied recursively to the newly aggregated network. This process continued until no further improvement in modularity was observed.

We adopted NetworkX (Hagberg et al. 2008), a Python library, to implement the Louvain method.

We summarized the five steps of the CoCoBin workflow in Algorithm 1, shown below.

#### Algorithm 1: CoCoBin

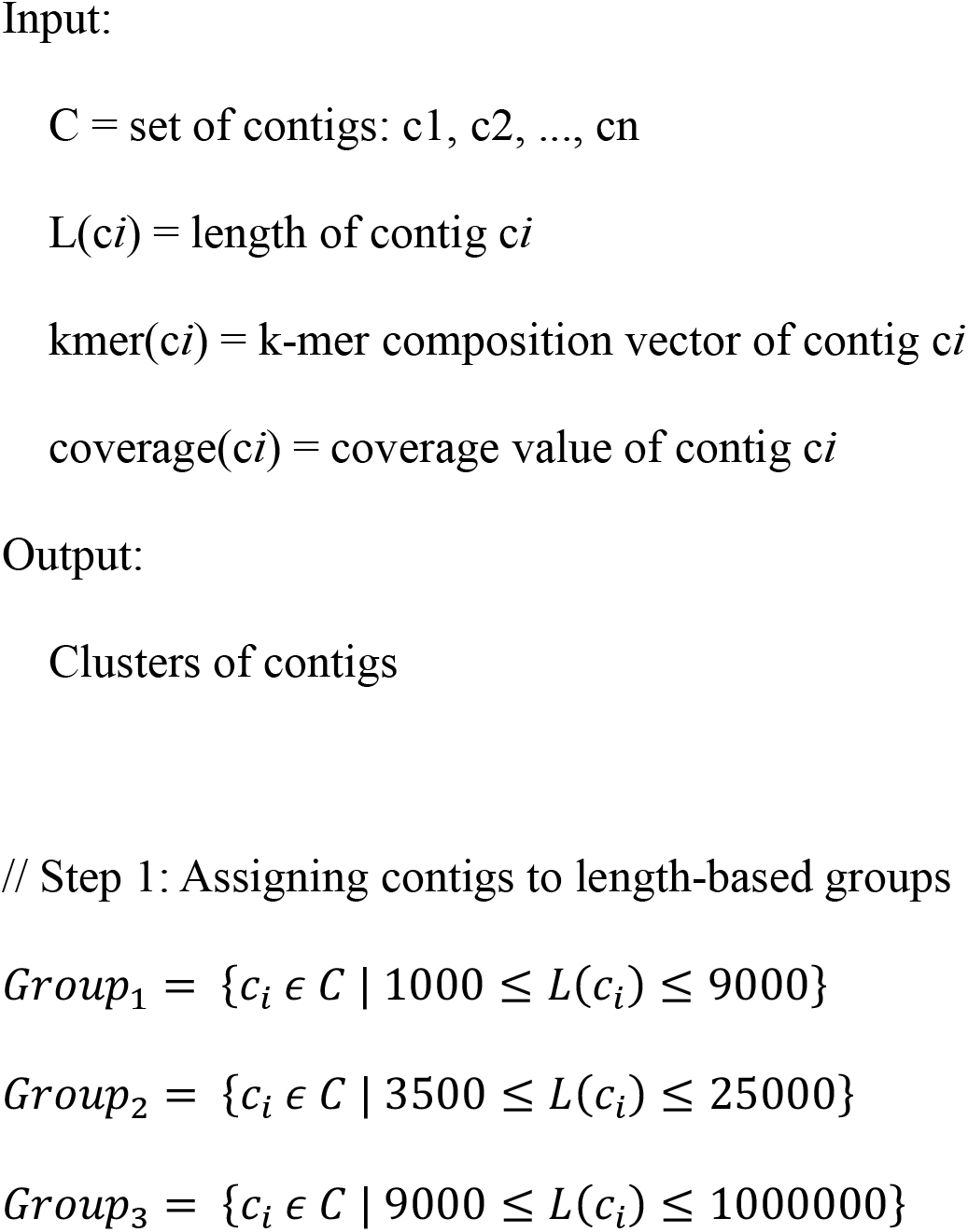
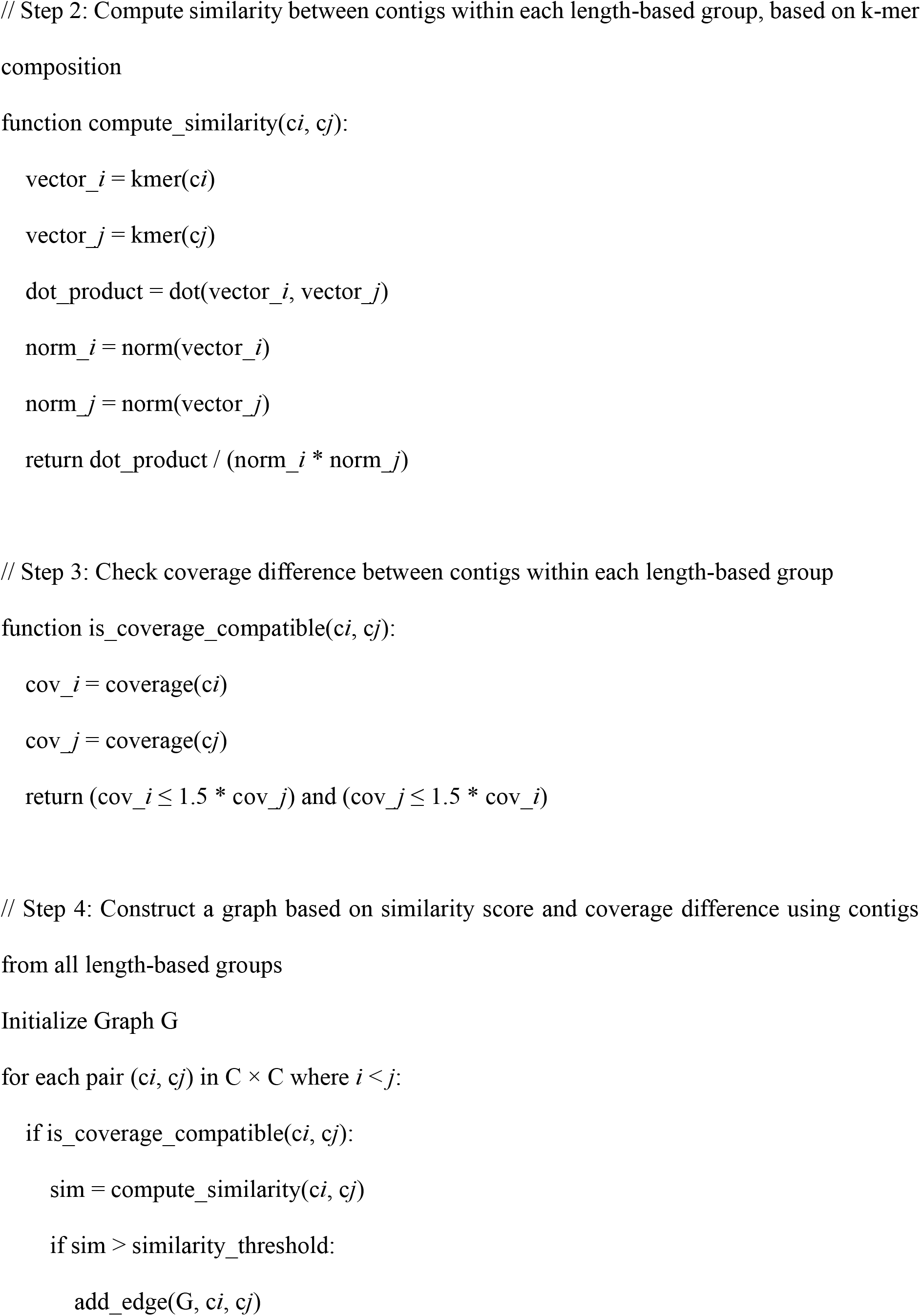
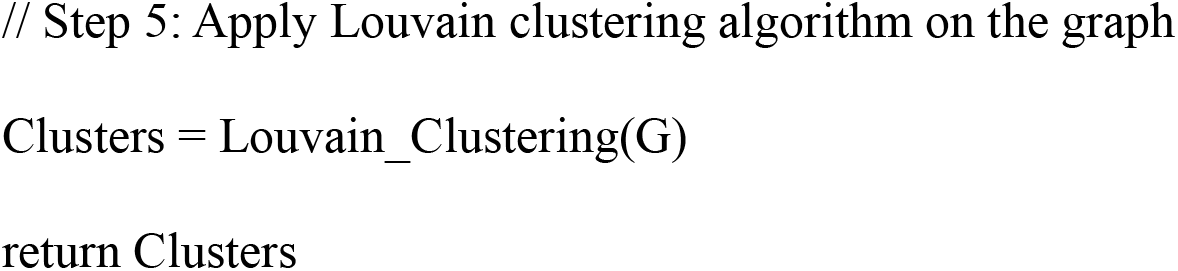

### 2.6 Datasets

To measure the generalization of CoCoBin and its effectiveness on metagenomic data binning, we obtained nine simulated datasets, five mock community datasets, and one real dataset from various sources, as presented below.

#### 2.6.1 Simulated Datasets

Sim-5G dataset (Xue et al. 2022) was simulated based on five species.

SRR8304764, SRR8304773, SRR8304775, and SRR8304776 datasets (Miller et al. 2019) were constructed from 51 bacterial strains isolated from the human gut. DNA was extracted from each isolate for SRR8304764 and SRR8304773, quantified, and mixed in equal molar amounts, while DNA was extracted from each isolate for SRR8304775 and SRR8304776, quantified, and mixed in staggered molar amounts. We obtained these raw read datasets from the Sequence Read Archive (SRA) at the National Center for Biotechnology Information (NCBI).

Short reads from the Sim79, Sim82, Sim92, and Sim98 datasets were generated using 79, 82, 92, and 98 bacterial species, respectively. We simulated the reads using InSilicoSeq (Gourlé et al. 2018), modeling a NovaSeq instrument with a mean read length of 151 bps. Details of the species used in each Sim dataset are provided in Supplementary Tables S20–S23.

#### 2.6.2 Mock Community Datasets

BMock12 dataset (NCBI Accession No. SRR8073716) (Sevim et al. 2019) comprise 12 bacterial species.

Amos HiLo datasets (NCBI Accession No. SRR11487931 and SRR11487935) (Amos et al. 2020) comprise 20 common gut microbes.

MBARC-26 dataset (NCBI Accession No. SRR3656745) (Singer et al. 2016) comprises 23 bacterial and three archaeal species.

SYNTH 64 dataset (NCBI Accession No. SRR606249) (Shakya et al. 2013) comprises 64 diverse bacterial and archaeal species.

#### 2.6.3 Real Dataset

The Sharon dataset (Sharon et al. 2013) comprises 18 runs from 11 fecal samples collected from the premature infant gut. The raw read dataset is accessible through SRA under accession number SRX144807.

### 2.7 Evaluation metrics

Since the reference genomes for the simulated and mock community datasets were available, BWA-MEM (Li 2013) was used to align the contigs to their corresponding reference genomes, allowing for the identification of the true species to which each contig belonged. Sharon is a real dataset. Hence, we used BLAST (Altschul et al. 1990) to search for matches in the NCBI nucleotide database. To facilitate a comprehensive evaluation, we converted the mappings derived from these contigs and reference genomes into a gold standard file format compatible with the Assessment of Metagenome BinnERs (AMBER) tool (Meyer et al. 2018). AMBER employed five key evaluation metrics to assess binning performance as follows. Accuracy measures the overall correctness of bin assignments. Purity indicates the proportion of correctly assigned contigs within each bin, reflecting precision. Completeness assesses recall by representing the proportion of contigs assigned to a bin relative to the total contigs in the reference genome. F1 score provides a harmonic mean of precision and recall, offering a balanced assessment of binning accuracy. The rand index evaluates clustering consistency by measuring the similarity between ground truth annotations and bin assignments. The rand index is a way to compare the similarity of results between two different clustering methods.

## 3. Results

To evaluate the effectiveness of the CoCoBin, we conducted a comprehensive comparison against several state-of-the-art clustering methods: BusyBee Web, CONCOCT (version 1.0.0), MaxBin 2.0 (version 2.2.7), MetaBAT 2 (version 2.15), and MetaDecoder (version v1.0.18). All tools were run with their default parameters to reconstruct genome clusters. The minimum contig length threshold in MetaDecoder was set to 2.5 Kb by default, which was consistent with the cutoff used by MetaBAT 2. In contrast, BusyBee Web, CONCOCT, and MaxBin 2 used a threshold of 1 Kb. We evaluated the genome clusters across various datasets, including nine simulated datasets, five mock community datasets, and one real dataset. The AMBER tool (version 2.0.4) was used to assess the results obtained from all binning tools. The outputs of the other tools were generated by executing each program with its respective default settings.

### 3.1 Results on simulated datasets

The evaluation results from multiple binning tools applied to simulated datasets demonstrate that CoCoBin outperforms other tools in terms of the number of bins identified across seven out of nine datasets (Fig. 3) —SRR8304764, SRR8304773, SRR8304775, SRR8304776, Sim79, Sim82, and Sim92 —and rank second on Sim98. Regarding F1 scores, which represent the balance between purity and completeness, our model ranks first in four out of nine datasets—SRR8304764, SRR8304773, SRR8304775, and SRR8304776 —and second in three datasets Sim79, Sim82, and Sim92. Fig. 4 illustrates the performance of the binning tools across various metrics, including accuracy, purity, completeness, F1 score, and rand index. The AMBER results for these datasets are provided in Supplementary Table S24.

**Fig. 3.**
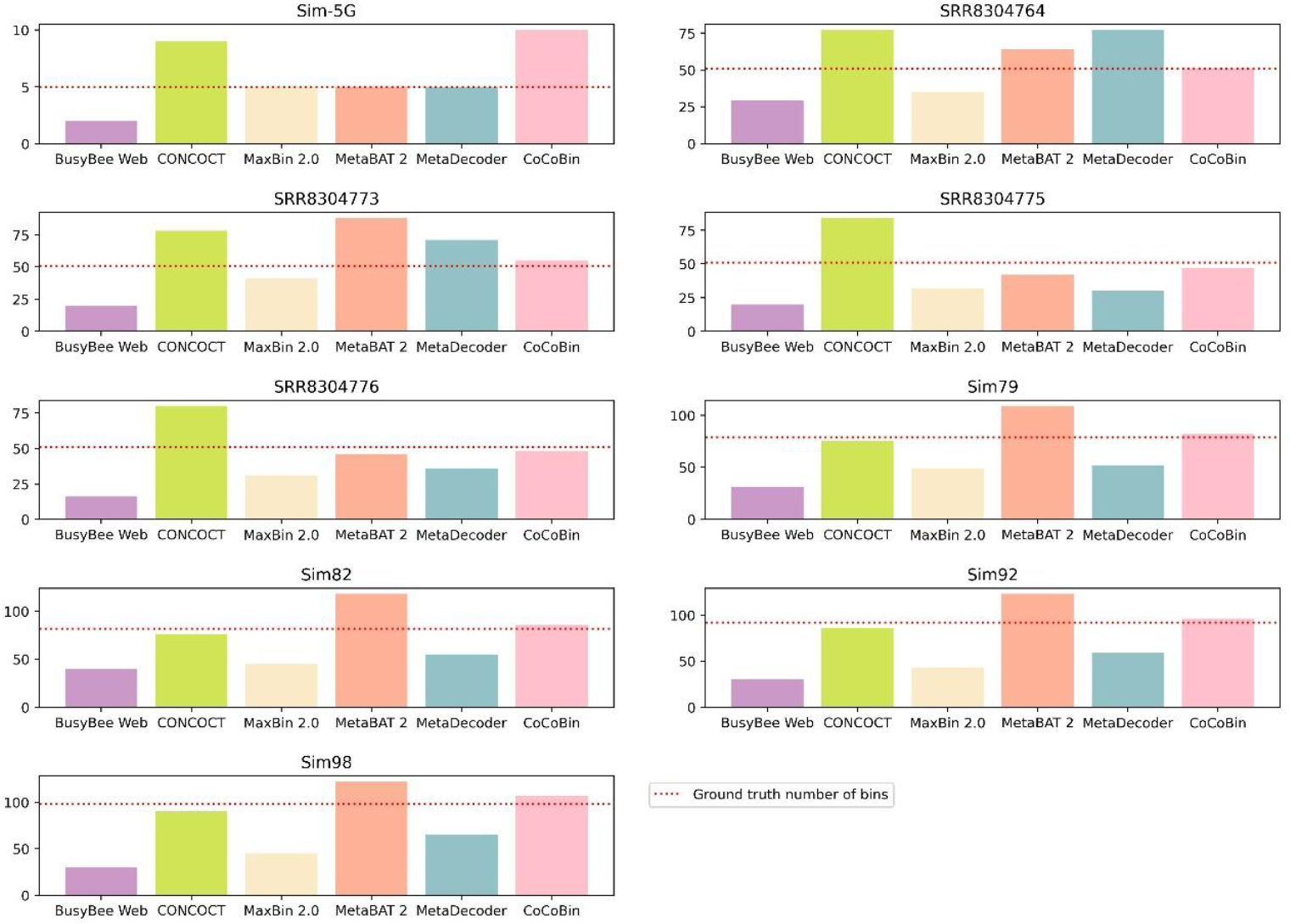
Number of bins identified by each tool compared with the ground truth number of bins (dotted line) on the nine simulated datasets.

**Fig. 4.**
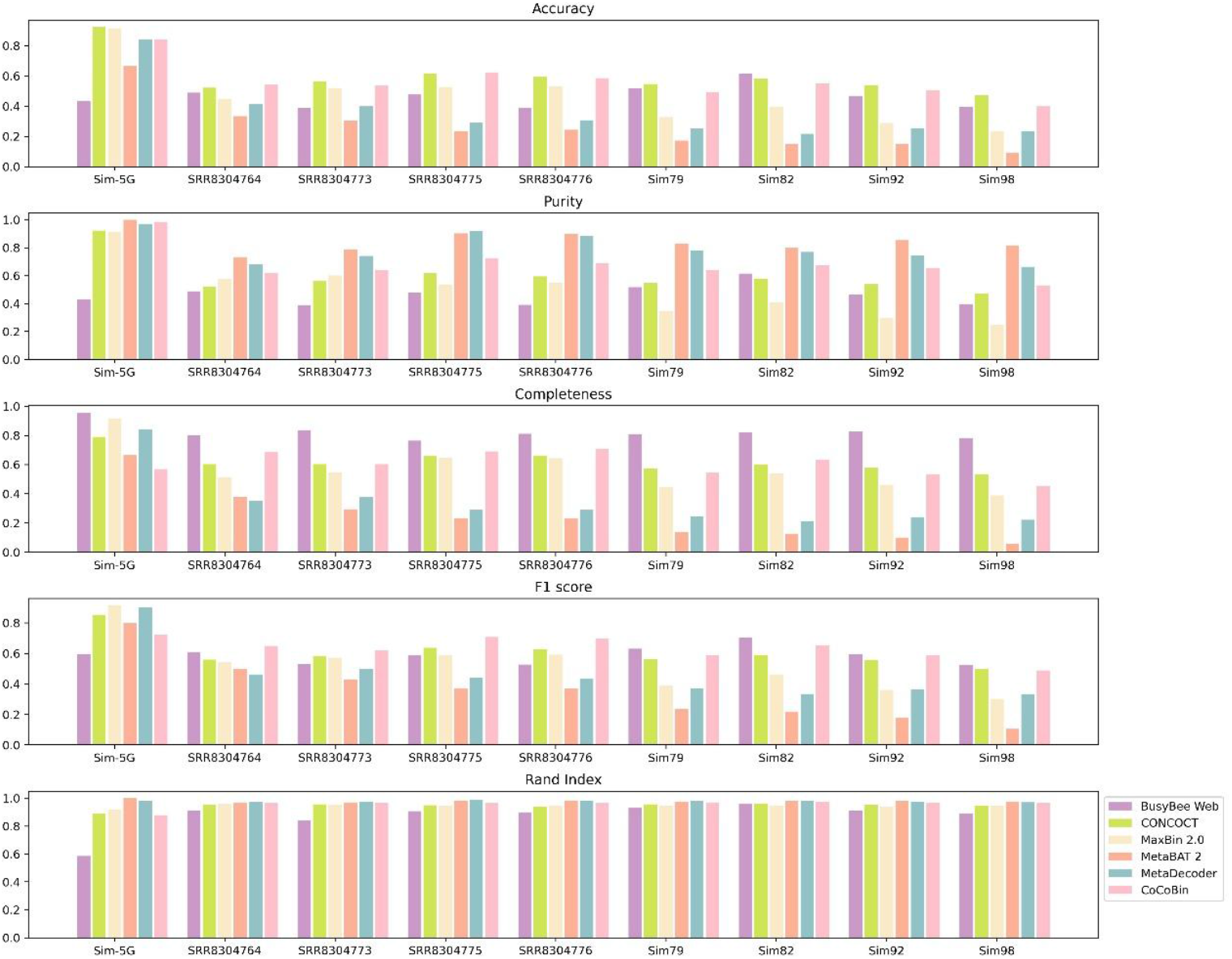
Comparison of binning performance metrics of the six binning tools on the nine simulated datasets.

### 3.2 Results on mock community datasets

The evaluation results from multiple binning tools applied to mock community datasets demonstrate that CoCoBin outperforms other tools in terms of the number of bins identified across three out of five datasets (Fig. 5) — SRR11487931, SRR11487935, and SRR606249 — and second on SRR3656745. In terms of F1 scores, our model ranks first on SRR3656745 and second on SRR11487931, SRR11487935, and SRR606249. Fig. 6 presents a comparative analysis of the binning tools’ performance metrics on the mock community datasets. The AMBER results for these datasets are provided in Supplementary Table S25.

**Fig. 5.**
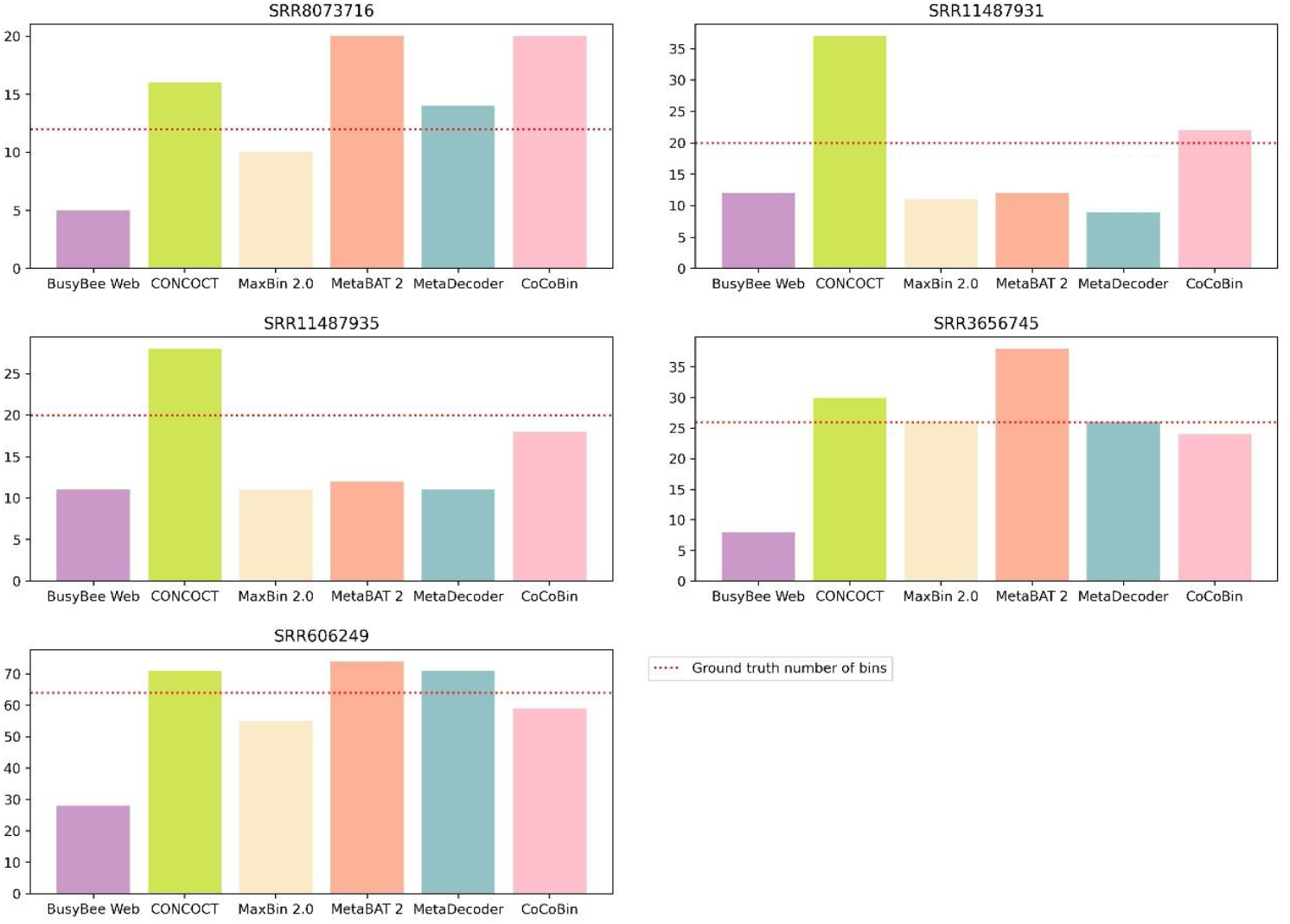
The number of bins identified by each tool compared with the ground truth number of bins (dotted line) on the five mock community datasets.

**Fig. 6.**
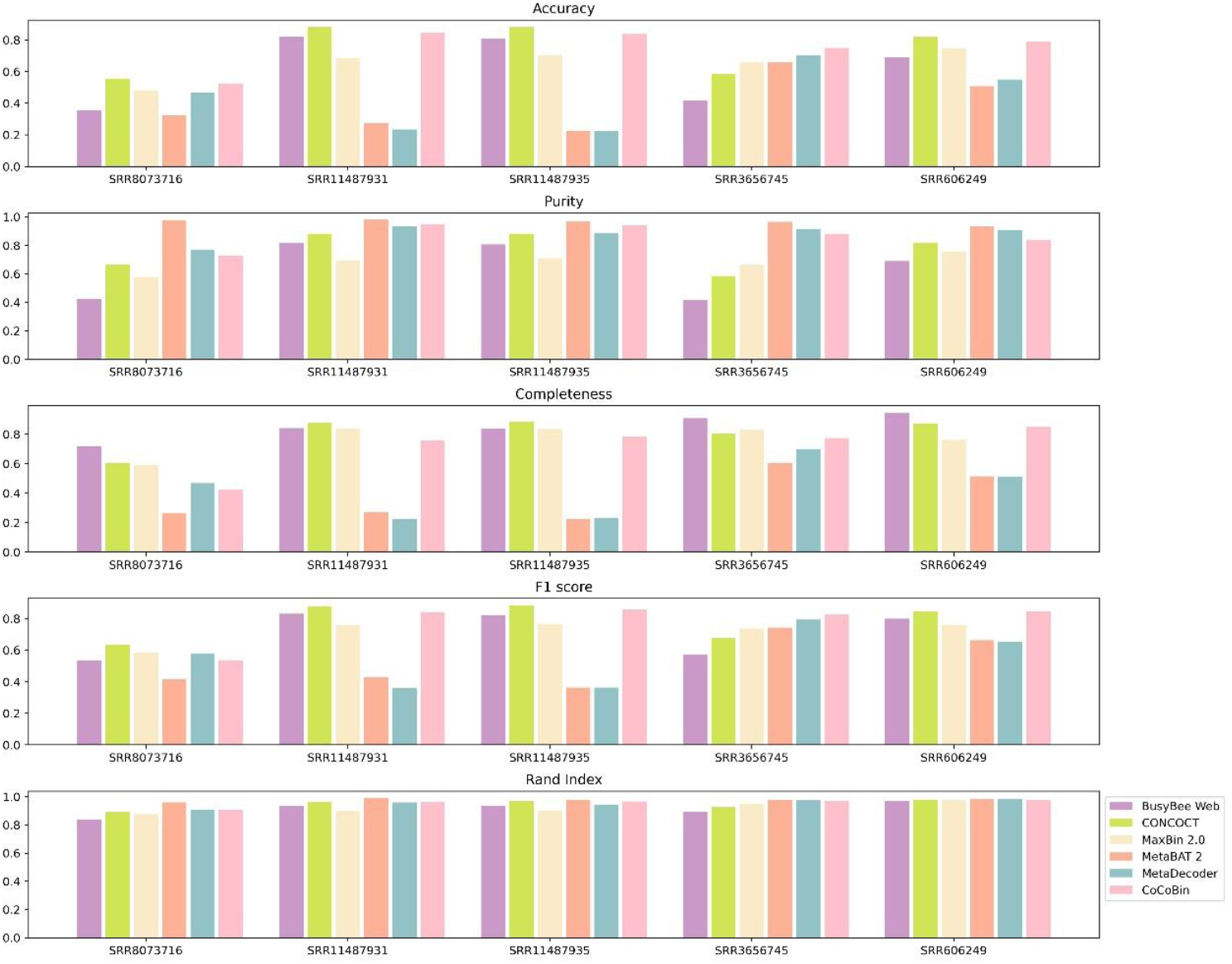
Comparison of binning performance metrics of the six binning tools on the five mock community datasets.

### 3.3 Results on real-world dataset

The evaluation results from the real-world Sharon dataset show that CoCoBin performs best compared to other tools in terms of the number of bins identified (Fig. 7), while ranking second in accuracy (0.799) and F1 score (0.690). Furthermore, Fig. 8 presents the performance of the binning tools across various metrics on the real-world dataset. The AMBER results for this dataset are provided in Supplementary Table S26.

**Fig. 7.**
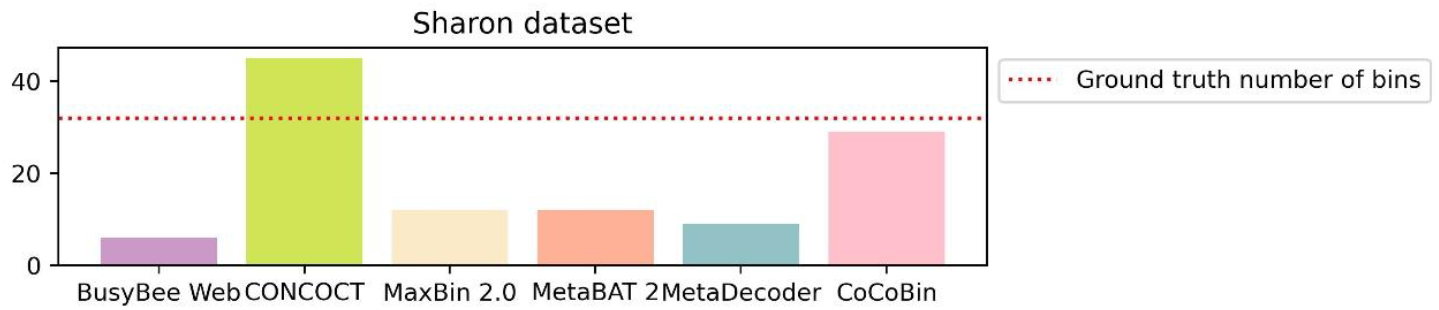
The number of bins identified by each tool compared with the ground truth number of bins (dotted line) on the real-world (Sharon) dataset.

**Fig. 8.**
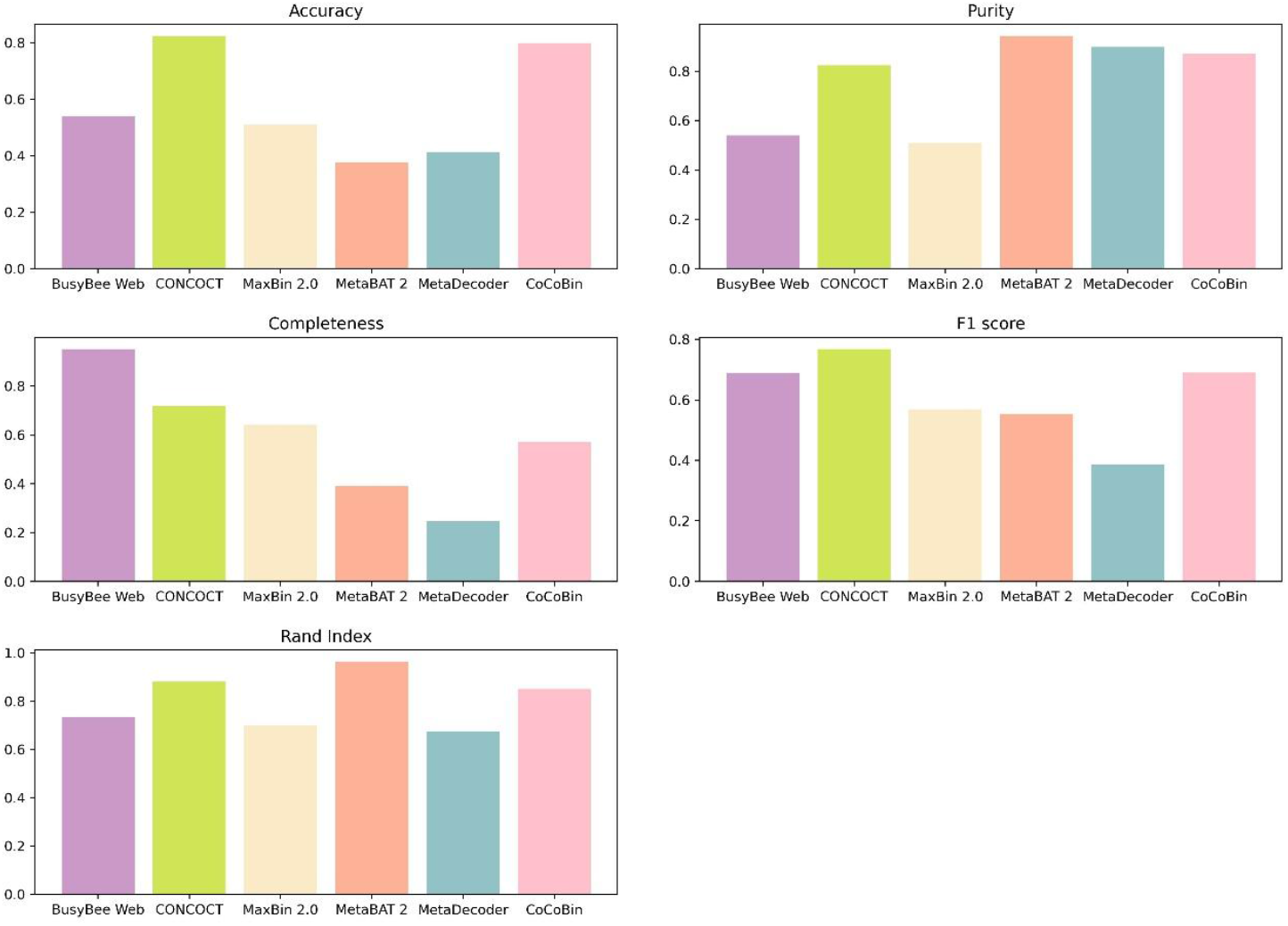
Comparison of binning performance metrics of the six binning tools on the real-world (Sharon) dataset.

## 4. Discussion and conclusion

The CoCoBin is a multi-stage binning method designed to enhance metagenomic analysis. It proceeds through the following stages: (1) assembling reads into contigs, (2) extracting compositional features, (3) computing contig similarity, (4) structuring complex networks, and (5) clustering. This study primarily focuses on the contig similarity computation stage, a critical component of the binning process. Contigs are initially grouped based on their length ranges. Similarity between contigs is then calculated using a combination of compositional similarity and coverage difference. This hybrid similarity metric, when integrated with the Louvain clustering algorithm, demonstrates strong performance in terms of the number of bins identified.

In simulated datasets, CoCoBin achieves outstanding results, ranking first in the number of bins identified in seven out of nine datasets and second in one. It also attains the highest F1 score in four datasets and ranks second in three. On mock community datasets, CoCoBin remains competitive, ranking first in the number of bins identified in three out of five datasets and second in one. It also achieves the highest F1 score in one dataset and ranks second in three. In evaluations using a real-world dataset, CoCoBin most accurately predicts the number of bins, coming closest to the actual number of species. However, its performance varies across other metrics: CONCOCT achieves the highest accuracy and F1 score, BusyBee Web leads in completeness, and MetaBAT 2 outperforms others in purity and Rand index. Despite these strengths, these tools are less accurate than CoCoBin in estimating the correct number of bins. From simulated, mock community, and real-world datasets, these outcomes suggest that CoCoBin is robust and generalizable across varying data types and complexities, demonstrating the model’s effectiveness in accurately clustering metagenomics data. However, within these clusters, a certain level of contamination between species persists, indicating the need for further refinement to minimize cross-species contamination. CoCoBin is not well suited for datasets with a small number of contigs, such as the Sim-5G and SRR8073716 datasets, which comprise only 248 and 903 contigs, respectively (see Supplementary Tables S1). In such cases, the sparsity of contigs within each group disrupts the underlying relationships and weakens the method’s effectiveness. The Sim98 dataset represents complex and highly diverse microbial communities characterized by high inter-species similarity. In such scenarios, this model still struggles to eliminate potential contamination due to the underlying complexity of the community structure.

In future research, we plan to develop a binning algorithm specifically designed for complex, highly diverse microbial communities with high inter-species similarity. This involves incorporating additional genomic features and adjusting parameter settings accordingly.

## Acknowledgements

We want to thank the department of computer engineering, faculty of engineering, Chulalongkorn university for providing computing facilities.

