## Supplementary Table S1-S26 for "CoCoBin: Graph-Based Metagenomic Binning via Composition–Coverage Separation"

**Supplementary Material for “CoCoBin”**

This supplementary material contains details of the experimental results from parameter tuning on simulated and mock community datasets, used to evaluate CoCoBin.

**Table S1** Summary of datasets with the number of contigs and the distribution of contigs’ length ranges used to evaluate CoCoBin

|  | **Dataset** | **Number of contigs** | | | |
| --- | --- | --- | --- | --- | --- |
|  |  | **1000 - 3500**  **bps** | **3501 – 10000 bps** | **10001 – 50000 bps** | **> 50000**  **bps** |
| Simulated datasets | Sim-5G | 47 | 33 | 64 | 104 |
|  | SRR8304764 | 4230 | 2403 | 2315 | 461 |
|  | SRR8304773 | 2434 | 1135 | 1544 | 562 |
|  | SRR8304775 | 8653 | 2421 | 1388 | 263 |
|  | SRR8304776 | 8516 | 2489 | 1370 | 269 |
|  | Sim79 | 13550 | 3497 | 1716 | 315 |
|  | Sim82 | 16601 | 3361 | 1693 | 334 |
|  | Sim92 | 12438 | 3342 | 1935 | 324 |
|  | Sim98 | 13321 | 3359 | 2107 | 347 |
| Mock community datasets | SRR8073716 | 304 | 173 | 225 | 201 |
|  | SRR11487931 | 4949 | 923 | 471 | 63 |
|  | SRR11487935 | 4819 | 1033 | 455 | 66 |
|  | SRR3656745 | 425 | 298 | 550 | 428 |
|  | SRR606249 | 5759 | 3500 | 2068 | 650 |

**Table S2** List of organisms with the number of contigs and the distribution of contigs’ length ranges of Sim-5G dataset

| **Organism Name** | **Number of contigs** | | | |
| --- | --- | --- | --- | --- |
|  | **1000 - 3500**  **bps** | **3501 – 10000**  **bps** | **10001 – 50000 bps** | **> 50000**  **bps** |
| Acetobacter pasteurianus | 18 | 15 | 16 | 20 |
| Aeromonas veronii | 4 | 1 | 13 | 23 |
| Amycolatopsis mediterranei | 5 | 3 | 6 | 32 |
| Arthrobacter arilaitensis | 20 | 14 | 29 | 20 |
| Azorhizobium caulinodans | 0 | 0 | 0 | 9 |

**Table S3** List of organisms with the number of contigs and the distribution of contigs’ length ranges of SRR8304764 dataset

| **Organism Name** | **Number of contigs** | | | |
| --- | --- | --- | --- | --- |
|  | **1000 - 3500**  **bps** | **3501 – 10000**  **bps** | **10001 – 50000 bps** | **> 50000**  **bps** |
| Alistipes indistinctus YIT 12060 | 1 | 2 | 5 | 5 |
| Bacteroides cellulosilyticus DSM 14838 | 24 | 19 | 18 | 2 |
| Bacteroides finegoldii DSM 17565 | 38 | 43 | 68 | 20 |
| Bacteroides intestinalis DSM 17393 | 20 | 33 | 35 | 26 |
| Bacteroides ovatus ATCC 8483 | 17 | 23 | 47 | 36 |
| Bacteroides plebeius DSM 17135 | 31 | 10 | 34 | 26 |
| Bacteroides stercoris ATCC 43183 | 22 | 22 | 52 | 18 |
| Bacteroides thetaiotaomicron 3731 | 66 | 60 | 46 | 4 |
| Bacteroides thetaiotaomicron 7330 | 67 | 58 | 56 | 2 |
| Bacteroides thetaiotaomicron VPI-5482 | 59 | 91 | 127 | 12 |
| Bacteroides uniformis ATCC 8492 | 20 | 12 | 21 | 16 |
| Bacteroides vulgatus ATCC 8482 | 107 | 163 | 135 | 4 |
| Bacteroidetes dorei DSM 17855 | 97 | 88 | 106 | 10 |
| Bifidobacterium adolescentis L2-32 | 83 | 80 | 83 | 0 |
| Bifidobacterium angulatum DSM 20098 | 68 | 52 | 58 | 5 |
| Bifidobacterium bifidum ATCC 29521 | 16 | 19 | 44 | 12 |
| Bifidobacterium dentium ATCC 27678 | 33 | 62 | 75 | 5 |
| Bifidobacterium pseudocatenulatum DSM 20438 | 136 | 114 | 50 | 3 |
| Blautia hansenii DSM 20583 | 94 | 61 | 80 | 4 |
| Blautia luti DSM 14534 | 63 | 40 | 50 | 20 |
| Citrobacter youngae ATCC 29220 | 7 | 8 | 12 | 24 |
| Clostridium asparagiforme DSM 15981 | 21 | 11 | 14 | 4 |
| Clostridium bolteae ATCC BAA-613 | 34 | 19 | 28 | 38 |
| Clostridium hathewayi DSM 13479 | 37 | 18 | 22 | 3 |
| Clostridium hylemonae DSM 15053 | 877 | 155 | 4 | 0 |
| Tyzzerella nexilis DSM 1787 | 107 | 66 | 53 | 6 |
| Clostridium ramosum DSM 1402 | 151 | 32 | 28 | 0 |
| Clostridium sp. M62/1 | 118 | 103 | 112 | 5 |
| Clostridium sporogenes ATCC 15579 | 90 | 26 | 23 | 0 |
| Clostridium symbiosum ATCC 14940 | 63 | 82 | 154 | 8 |
| Collinsella intestinalis DSM 13280 | 4 | 5 | 10 | 9 |
| Collinsella stercoris DSM 13279 | 36 | 71 | 59 | 1 |
| Coprococcus comes ATCC 27758 | 32 | 12 | 7 | 0 |
| Dorea formicigenerans ATCC 27755 | 144 | 39 | 8 | 0 |
| Edwardsiella tarda ATCC 23685 | 5 | 2 | 15 | 7 |
| Enterobacter cancerogenus ATCC 35316 | 1 | 4 | 4 | 21 |
| Escherichia fergusonii ATCC 35469 | 22 | 11 | 29 | 30 |
| Eubacterium biforme DSM 3989 | 106 | 29 | 9 | 0 |
| Holdemania filiformis DSM 12042 | 22 | 8 | 30 | 4 |
| Eubacterium eligens ATCC 27750 | 108 | 52 | 75 | 3 |
| Lactobacillus reuteri DSM 20016 | 386 | 96 | 6 | 0 |
| Lactobacillus ruminis DSM 20403 | 106 | 51 | 10 | 0 |
| Marvinbryantia formatexigens DSM 14469 | 115 | 111 | 116 | 12 |
| Megamonas funiformis YIT 11815 | 48 | 37 | 45 | 8 |
| Parabacteroides johnsonii DSM 18315 | 8 | 18 | 21 | 2 |
| Parabacteroides merdae ATCC 43184 | 18 | 19 | 41 | 10 |
| Proteus penneri ATCC 35198 | 44 | 7 | 0 | 0 |
| Roseburia intestinalis L1-82 | 266 | 131 | 88 | 2 |
| Ruminococcus gnavus ATCC 29149 | 45 | 21 | 27 | 20 |
| Streptococcus infantarius ATCC BAA-102 | 113 | 93 | 47 | 0 |
| Subdoligranulum variabile DSM 15176 | 34 | 14 | 28 | 14 |

**Table S4** List of organisms with the number of contigs and the distribution of contigs’ length ranges of SRR8304773 dataset

| **Organism Name** | **Number of contigs** | | | |
| --- | --- | --- | --- | --- |
|  | **1000 - 3500**  **bps** | **3501 – 10000**  **bps** | **10001 – 50000 bps** | **> 50000**  **bps** |
| Alistipes indistinctus YIT 12060 | 2 | 0 | 3 | 5 |
| Bacteroides cellulosilyticus DSM 14838 | 11 | 18 | 12 | 3 |
| Bacteroides finegoldii DSM 17565 | 37 | 34 | 55 | 16 |
| Bacteroides intestinalis DSM 17393 | 15 | 16 | 21 | 12 |
| Bacteroides ovatus ATCC 8483 | 20 | 21 | 51 | 33 |
| Bacteroides plebeius DSM 17135 | 22 | 7 | 18 | 27 |
| Bacteroides stercoris ATCC 43183 | 26 | 14 | 45 | 18 |
| Bacteroides thetaiotaomicron 3731 | 65 | 61 | 43 | 6 |
| Bacteroides thetaiotaomicron 7330 | 63 | 60 | 52 | 5 |
| Bacteroides thetaiotaomicron VPI-5482 | 72 | 89 | 124 | 13 |
| Bacteroides uniformis ATCC 8492 | 16 | 9 | 15 | 18 |
| Bacteroides vulgatus ATCC 8482 | 49 | 99 | 107 | 9 |
| Bacteroidetes dorei DSM 17855 | 45 | 40 | 79 | 13 |
| Bifidobacterium adolescentis L2-32 | 21 | 10 | 31 | 15 |
| Bifidobacterium angulatum DSM 20098 | 7 | 7 | 18 | 11 |
| Bifidobacterium bifidum ATCC 29521 | 3 | 4 | 4 | 14 |
| Bifidobacterium dentium ATCC 27678 | 9 | 4 | 6 | 19 |
| Bifidobacterium pseudocatenulatum DSM 20438 | 18 | 26 | 28 | 13 |
| Blautia hansenii DSM 20583 | 37 | 13 | 21 | 18 |
| Blautia luti DSM 14534 | 25 | 8 | 16 | 25 |
| Citrobacter youngae ATCC 29220 | 1 | 6 | 7 | 18 |
| Clostridium asparagiforme DSM 15981 | 7 | 4 | 7 | 2 |
| Clostridium bolteae ATCC BAA-613 | 17 | 16 | 27 | 34 |
| Clostridium hathewayi DSM 13479 | 18 | 12 | 14 | 2 |
| Clostridium hylemonae DSM 15053 | 60 | 88 | 102 | 13 |
| Tyzzerella nexilis DSM 1787 | 55 | 33 | 36 | 10 |
| Clostridium ramosum DSM 1402 | 309 | 47 | 47 | 2 |
| Clostridium sp. M62/1 | 25 | 15 | 22 | 18 |
| Clostridium sporogenes ATCC 15579 | 495 | 31 | 33 | 0 |
| Clostridium symbiosum ATCC 14940 | 36 | 48 | 124 | 12 |
| Collinsella intestinalis DSM 13280 | 5 | 2 | 9 | 5 |
| Collinsella stercoris DSM 13279 | 10 | 11 | 20 | 4 |
| Coprococcus comes ATCC 27758 | 17 | 6 | 1 | 0 |
| Dorea formicigenerans ATCC 27755 | 331 | 59 | 34 | 0 |
| Edwardsiella tarda ATCC 23685 | 5 | 5 | 10 | 6 |
| Enterobacter cancerogenus ATCC 35316 | 2 | 2 | 4 | 14 |
| Escherichia fergusonii ATCC 35469 | 6 | 8 | 10 | 19 |
| Eubacterium biforme DSM 3989 | 131 | 29 | 26 | 0 |
| Holdemania filiformis DSM 12042 | 31 | 10 | 18 | 4 |
| Eubacterium eligens ATCC 27750 | 41 | 15 | 32 | 20 |
| Lactobacillus reuteri DSM 20016 | 30 | 34 | 40 | 7 |
| Lactobacillus ruminis DSM 20403 | 37 | 19 | 9 | 0 |
| Marvinbryantia formatexigens DSM 14469 | 15 | 7 | 27 | 26 |
| Megamonas funiformis YIT 11815 | 18 | 5 | 24 | 10 |
| Parabacteroides johnsonii DSM 18315 | 13 | 9 | 14 | 0 |
| Parabacteroides merdae ATCC 43184 | 27 | 12 | 25 | 11 |
| Proteus penneri ATCC 35198 | 32 | 5 | 0 | 0 |
| Roseburia intestinalis L1-82 | 47 | 24 | 40 | 6 |
| Ruminococcus gnavus ATCC 29149 | 31 | 20 | 21 | 19 |
| Streptococcus infantarius ATCC BAA-102 | 6 | 1 | 5 | 2 |
| Subdoligranulum variabile DSM 15176 | 13 | 12 | 7 | 5 |

**Table S5** List of organisms with the number of contigs and the distribution of contigs’ length ranges of SRR8304775 dataset

| **Organism Name** | **Number of contigs** | | | |
| --- | --- | --- | --- | --- |
|  | **1000 - 3500**  **bps** | **3501 – 10000**  **bps** | **10001 – 50000 bps** | **> 50000**  **bps** |
| Alistipes indistinctus YIT 12060 | 2 | 0 | 3 | 5 |
| Bacteroides cellulosilyticus DSM 14838 | 11 | 18 | 12 | 3 |
| Bacteroides finegoldii DSM 17565 | 37 | 34 | 55 | 16 |
| Bacteroides intestinalis DSM 17393 | 15 | 16 | 21 | 12 |
| Bacteroides ovatus ATCC 8483 | 20 | 21 | 51 | 33 |
| Bacteroides plebeius DSM 17135 | 22 | 7 | 18 | 27 |
| Bacteroides stercoris ATCC 43183 | 26 | 14 | 45 | 18 |
| Bacteroides thetaiotaomicron 3731 | 65 | 61 | 43 | 6 |
| Bacteroides thetaiotaomicron 7330 | 63 | 60 | 52 | 5 |
| Bacteroides thetaiotaomicron VPI-5482 | 72 | 89 | 124 | 13 |
| Bacteroides uniformis ATCC 8492 | 16 | 9 | 15 | 18 |
| Bacteroides vulgatus ATCC 8482 | 49 | 99 | 107 | 9 |
| Bacteroidetes dorei DSM 17855 | 45 | 40 | 79 | 13 |
| Bifidobacterium adolescentis L2-32 | 21 | 10 | 31 | 15 |
| Bifidobacterium angulatum DSM 20098 | 7 | 7 | 18 | 11 |
| Bifidobacterium bifidum ATCC 29521 | 3 | 4 | 4 | 14 |
| Bifidobacterium dentium ATCC 27678 | 9 | 4 | 6 | 19 |
| Bifidobacterium pseudocatenulatum DSM 20438 | 18 | 26 | 28 | 13 |
| Blautia hansenii DSM 20583 | 37 | 13 | 21 | 18 |
| Blautia luti DSM 14534 | 25 | 8 | 16 | 25 |
| Citrobacter youngae ATCC 29220 | 1 | 6 | 7 | 18 |
| Clostridium asparagiforme DSM 15981 | 7 | 4 | 7 | 2 |
| Clostridium bolteae ATCC BAA-613 | 17 | 16 | 27 | 34 |
| Clostridium hathewayi DSM 13479 | 18 | 12 | 14 | 2 |
| Clostridium hylemonae DSM 15053 | 60 | 88 | 102 | 13 |
| Tyzzerella nexilis DSM 1787 | 55 | 33 | 36 | 10 |
| Clostridium ramosum DSM 1402 | 309 | 47 | 47 | 2 |
| Clostridium sp. M62/1 | 25 | 15 | 22 | 18 |
| Clostridium sporogenes ATCC 15579 | 495 | 31 | 33 | 0 |
| Clostridium symbiosum ATCC 14940 | 36 | 48 | 124 | 12 |
| Collinsella intestinalis DSM 13280 | 5 | 2 | 9 | 5 |
| Collinsella stercoris DSM 13279 | 10 | 11 | 20 | 4 |
| Coprococcus comes ATCC 27758 | 17 | 6 | 1 | 0 |
| Dorea formicigenerans ATCC 27755 | 331 | 59 | 34 | 0 |
| Edwardsiella tarda ATCC 23685 | 5 | 5 | 10 | 6 |
| Enterobacter cancerogenus ATCC 35316 | 2 | 2 | 4 | 14 |
| Escherichia fergusonii ATCC 35469 | 6 | 8 | 10 | 19 |
| Eubacterium biforme DSM 3989 | 131 | 29 | 26 | 0 |
| Holdemania filiformis DSM 12042 | 31 | 10 | 18 | 4 |
| Eubacterium eligens ATCC 27750 | 41 | 15 | 32 | 20 |
| Lactobacillus reuteri DSM 20016 | 30 | 34 | 40 | 7 |
| Lactobacillus ruminis DSM 20403 | 37 | 19 | 9 | 0 |
| Marvinbryantia formatexigens DSM 14469 | 15 | 7 | 27 | 26 |
| Megamonas funiformis YIT 11815 | 18 | 5 | 24 | 10 |
| Parabacteroides johnsonii DSM 18315 | 13 | 9 | 14 | 0 |
| Parabacteroides merdae ATCC 43184 | 27 | 12 | 25 | 11 |
| Proteus penneri ATCC 35198 | 32 | 5 | 0 | 0 |
| Roseburia intestinalis L1-82 | 47 | 24 | 40 | 6 |
| Ruminococcus gnavus ATCC 29149 | 31 | 20 | 21 | 19 |
| Streptococcus infantarius ATCC BAA-102 | 6 | 1 | 5 | 2 |
| Subdoligranulum variabile DSM 15176 | 13 | 12 | 7 | 5 |

**Table S6** List of organisms with the number of contigs and the distribution of contigs’ length ranges of SRR8304776 dataset

| **Organism Name** | **Number of contigs** | | | |
| --- | --- | --- | --- | --- |
|  | **1000 - 3500**  **bps** | **3501 – 10000**  **bps** | **10001 – 50000 bps** | **> 50000**  **bps** |
| Alistipes indistinctus YIT 12060 | 181 | 0 | 0 | 0 |
| Bacteroides cellulosilyticus DSM 14838 | 18 | 18 | 8 | 4 |
| Bacteroides finegoldii DSM 17565 | 101 | 78 | 111 | 6 |
| Bacteroides intestinalis DSM 17393 | 33 | 32 | 51 | 16 |
| Bacteroides ovatus ATCC 8483 | 79 | 65 | 93 | 37 |
| Bacteroides plebeius DSM 17135 | 1 | 1 | 1 | 0 |
| Bacteroides stercoris ATCC 43183 | 20 | 15 | 43 | 19 |
| Bacteroides thetaiotaomicron 3731 | 53 | 49 | 30 | 8 |
| Bacteroides thetaiotaomicron 7330 | 93 | 127 | 160 | 20 |
| Bacteroides thetaiotaomicron VPI-5482 | 27 | 28 | 29 | 3 |
| Bacteroides uniformis ATCC 8492 | 16 | 15 | 18 | 14 |
| Bacteroides vulgatus ATCC 8482 | 16 | 15 | 45 | 30 |
| Bacteroidetes dorei DSM 17855 | 493 | 56 | 1 | 0 |
| Bifidobacterium adolescentis L2-32 | 2 | 0 | 0 | 0 |
| Bifidobacterium angulatum DSM 20098 | 36 | 54 | 61 | 1 |
| Bifidobacterium bifidum ATCC 29521 | 2 | 0 | 0 | 0 |
| Bifidobacterium dentium ATCC 27678 | 337 | 199 | 34 | 0 |
| Bifidobacterium pseudocatenulatum DSM 20438 | 369 | 18 | 1 | 0 |
| Blautia hansenii DSM 20583 | 238 | 9 | 0 | 0 |
| Blautia luti DSM 14534 | 125 | 106 | 68 | 15 |
| Citrobacter youngae ATCC 29220 | 2 | 7 | 14 | 20 |
| Clostridium asparagiforme DSM 15981 | 17 | 7 | 10 | 5 |
| Clostridium bolteae ATCC BAA-613 | 1194 | 421 | 44 | 0 |
| Clostridium hathewayi DSM 13479 | 73 | 71 | 45 | 4 |
| Clostridium hylemonae DSM 15053 | 76 | 0 | 0 | 0 |
| Tyzzerella nexilis DSM 1787 | 43 | 0 | 1 | 0 |
| Clostridium ramosum DSM 1402 | 93 | 30 | 12 | 0 |
| Clostridium sp. M62/1 | 47 | 34 | 51 | 18 |
| Clostridium sporogenes ATCC 15579 | 53 | 28 | 2 | 0 |
| Clostridium symbiosum ATCC 14940 | 220 | 295 | 131 | 0 |
| Collinsella intestinalis DSM 13280 | 94 | 126 | 44 | 0 |
| Collinsella stercoris DSM 13279 | 269 | 0 | 0 | 0 |
| Coprococcus comes ATCC 27758 | 404 | 10 | 0 | 0 |
| Dorea formicigenerans ATCC 27755 | 152 | 35 | 5 | 0 |
| Edwardsiella tarda ATCC 23685 | 232 | 0 | 0 | 0 |
| Enterobacter cancerogenus ATCC 35316 | 2 | 2 | 2 | 2 |
| Escherichia fergusonii ATCC 35469 | 1067 | 154 | 4 | 0 |
| Eubacterium biforme DSM 3989 | 6 | 1 | 0 | 0 |
| Holdemania filiformis DSM 12042 | 169 | 2 | 1 | 0 |
| Eubacterium eligens ATCC 27750 | 230 | 120 | 33 | 0 |
| Lactobacillus reuteri DSM 20016 | 1 | 0 | 0 | 0 |
| Lactobacillus ruminis DSM 20403 | 158 | 49 | 4 | 0 |
| Marvinbryantia formatexigens DSM 14469 | 25 | 33 | 72 | 26 |
| Megamonas funiformis YIT 11815 | 29 | 0 | 0 | 0 |
| Parabacteroides johnsonii DSM 18315 | 29 | 28 | 26 | 3 |
| Parabacteroides merdae ATCC 43184 | 20 | 28 | 43 | 7 |
| Proteus penneri ATCC 35198 | 221 | 2 | 0 | 0 |
| Roseburia intestinalis L1-82 | 67 | 35 | 56 | 8 |
| Ruminococcus gnavus ATCC 29149 | 514 | 9 | 0 | 0 |
| Streptococcus infantarius ATCC BAA-102 | 12 | 11 | 15 | 3 |
| Subdoligranulum variabile DSM 15176 | 757 | 66 | 1 | 0 |

**Table S7** List of organisms with the number of contigs and the distribution of contigs’ length ranges of Sim79 dataset

| **Organism Name** | **Number of contigs** | | | |
| --- | --- | --- | --- | --- |
|  | **1000 - 3500**  **bps** | **3501 – 10000**  **bps** | **10001 – 50000 bps** | **> 50000**  **bps** |
| [Clostridium] asparagiforme DSM 15981 | 635 | 73 | 57 | 5 |
| [Clostridium] hylemonae DSM 15053 | 469 | 10 | 7 | 4 |
| [Clostridium] nexile DSM 1787 | 225 | 132 | 65 | 10 |
| [Clostridium] symbiosum ATCC 14940 | 49 | 85 | 157 | 12 |
| Acidithiobacillus ferrivorans SS3 | 379 | 3 | 1 | 0 |
| Acidovorax citrulli AAC00-1 | 37 | 1 | 0 | 0 |
| Actinobacillus succinogenes 130Z | 519 | 24 | 0 | 0 |
| Akkermansia muciniphila ATCC BAA-835 | 275 | 226 | 44 | 0 |
| Alistipes indistinctus YIT 12060 | 291 | 30 | 0 | 0 |
| Aminobacterium colombiense DSM 12261 | 86 | 0 | 0 | 0 |
| Arcobacter nitrofigilis DSM 7299 | 7 | 0 | 0 | 0 |
| Bacillus cellulosilyticus DSM 2522 | 359 | 1 | 0 | 0 |
| Bacillus cytotoxicus NVH 391-98 | 0 | 2 | 0 | 0 |
| Bacteroides cellulosilyticus DSM 14838 | 290 | 44 | 18 | 1 |
| Bacteroides finegoldii DSM 17565 | 164 | 108 | 69 | 11 |
| Bacteroides intestinalis DSM 17393 | 81 | 3 | 12 | 2 |
| Bacteroides ovatus strain ATCC 8483 | 2 | 0 | 0 | 0 |
| Bacteroides plebeius DSM 17135 | 85 | 44 | 37 | 11 |
| Bacteroides stercoris ATCC 43183 | 293 | 154 | 63 | 6 |
| Bacteroides thetaiotaomicron 3731 | 263 | 87 | 58 | 31 |
| Bacteroides thetaiotaomicron 7330 | 20 | 1 | 6 | 3 |
| Bacteroides thetaiotaomicron VPI-5482 | 13 | 2 | 4 | 8 |
| Bacteroides uniformis ATCC 8492 | 181 | 57 | 36 | 14 |
| Bacteroides vulgatus ATCC 8482 | 3 | 0 | 0 | 0 |
| Beijerinckia indica subsp. indica ATCC 9039 | 3 | 3 | 4 | 1 |
| Bifidobacterium adolescentis L2-32 | 395 | 61 | 5 | 5 |
| Bifidobacterium angulatum DSM 20098 | 3 | 0 | 0 | 0 |
| Bifidobacterium bifidum ATCC 29521 | 56 | 1 | 0 | 0 |
| Bifidobacterium dentium ATCC 27678 | 379 | 9 | 0 | 0 |
| Bifidobacterium pseudocatenulatum DSM 20438 | 289 | 134 | 18 | 4 |
| Blautia hansenii DSM 20583 | 10 | 2 | 0 | 0 |
| Blautia luti DSM 14534 | 103 | 35 | 41 | 18 |
| Brevundimonas subvibrioides ATCC 15264 | 2 | 1 | 0 | 0 |
| Burkholderia orbicola MC0-3 | 307 | 42 | 0 | 0 |
| Calothrix sp. PCC 6303 | 6 | 0 | 1 | 3 |
| Caulobacter segnis ATCC 21756 | 7 | 0 | 0 | 0 |
| Chlamydia trachomatis G/9301 | 236 | 69 | 3 | 0 |
| Chlorobium phaeobacteroides BS1 | 28 | 0 | 0 | 0 |
| Chromohalobacter salexigens DSM 3043 | 558 | 10 | 0 | 0 |
| Citrobacter youngae ATCC 29220 | 118 | 4 | 4 | 5 |
| Clostridium bolteae ATCC BAA-613 | 0 | 1 | 3 | 1 |
| Clostridium lentocellum DSM 5427 | 1 | 1 | 0 | 1 |
| Clostridium sp. M62/1 | 454 | 88 | 28 | 10 |
| Clostridium sporogenes ATCC 15579 | 475 | 300 | 71 | 0 |
| Collinsella intestinalis DSM 13280 | 104 | 30 | 1 | 0 |
| Collinsella stercoris DSM 13279 | 165 | 13 | 19 | 1 |
| Coprococcus comes ATCC 27758 | 122 | 14 | 24 | 5 |
| Cryptobacterium curtum DSM 15641 | 40 | 0 | 0 | 0 |
| Cyanobacterium aponinum PCC 10605 | 45 | 2 | 1 | 1 |
| Cyanobium gracile PCC 6307 | 9 | 1 | 0 | 0 |
| Dehalococcoides mccartyi BAV1 | 260 | 5 | 0 | 0 |
| Desulfosporosinus acidiphilus SJ4 | 177 | 2 | 0 | 1 |
| Desulfotomaculum acetoxidans DSM 771 | 2 | 0 | 0 | 0 |
| Desulfovibrio vulgaris str. 'Miyazaki F' | 23 | 0 | 1 | 0 |
| Desulfurispirillum indicum S5 | 5 | 1 | 0 | 0 |
| Dorea formicigenerans ATCC 27755 | 140 | 74 | 67 | 7 |
| Edwardsiella tarda ATCC 23685 | 101 | 42 | 32 | 1 |
| Eggerthella lenta DSM 2243 | 2 | 0 | 1 | 0 |
| Enterobacter cancerogenus ATCC 35316 | 39 | 4 | 3 | 3 |
| Escherichia fergusonii ATCC 35469 | 16 | 3 | 1 | 0 |
| Fervidobacterium nodosum Rt17-B1 | 437 | 121 | 2 | 0 |
| Fluviicola taffensis DSM 16823 | 5 | 3 | 0 | 0 |
| Holdemanella biformis DSM 3989 | 243 | 94 | 32 | 3 |
| Holdemania filiformis DSM 12042 | 420 | 139 | 50 | 3 |
| Hungatella hathewayi DSM 13479 | 441 | 330 | 167 | 8 |
| Lachnospira eligens ATCC 27750 | 12 | 22 | 25 | 0 |
| Lactobacillus reuteri DSM 20016 | 131 | 0 | 0 | 0 |
| Ligilactobacillus ruminis DSM 20403 = NBRC 102161 | 16 | 34 | 51 | 8 |
| Marvinbryantia formatexigens DSM 14469 | 98 | 27 | 51 | 23 |
| Megamonas funiformis YIT 11815 | 134 | 124 | 56 | 4 |
| Parabacteroides johnsonii DSM 18315 | 337 | 85 | 30 | 1 |
| Parabacteroides merdae ATCC 43184 | 319 | 93 | 79 | 12 |
| Phocaeicola dorei DSM 17855 | 666 | 62 | 21 | 10 |
| Proteus penneri ATCC 35198 | 503 | 125 | 5 | 0 |
| Roseburia intestinalis L1-82 | 175 | 111 | 103 | 9 |
| Ruminococcus gnavus ATCC 29149 | 134 | 31 | 22 | 22 |
| Streptococcus infantarius ATCC BAA-102 | 10 | 21 | 30 | 10 |
| Subdoligranulum variabile DSM 15176 | 15 | 19 | 25 | 1 |
| Thomasclavelia ramosa DSM 1402 | 48 | 17 | 5 | 16 |

**Table S8** List of organisms with the number of contigs and the distribution of contigs’ length ranges of Sim82 dataset

| **Organism Name** | **Number of contigs** | | | |
| --- | --- | --- | --- | --- |
|  | **1000 - 3500**  **bps** | **3501 – 10000**  **bps** | **10001 – 50000 bps** | **> 50000**  **bps** |
| [Clostridium] asparagiforme DSM 15981 | 197 | 112 | 61 | 9 |
| [Clostridium] hylemonae DSM 15053 | 73 | 27 | 18 | 0 |
| [Clostridium] nexile DSM 1787 | 184 | 73 | 39 | 6 |
| [Clostridium] symbiosum ATCC 14940 | 84 | 79 | 151 | 9 |
| Alistipes indistinctus YIT 12060 | 153 | 26 | 46 | 0 |
| Bacteroides cellulosilyticus DSM 14838 | 674 | 122 | 60 | 1 |
| Bacteroides finegoldii DSM 17565 | 294 | 86 | 64 | 15 |
| Bacteroides intestinalis DSM 17393 | 126 | 55 | 8 | 1 |
| Bacteroides plebeius DSM 17135 | 227 | 47 | 22 | 16 |
| Bacteroides stercoris ATCC 43183 | 192 | 38 | 32 | 13 |
| Bacteroides thetaiotaomicron 3731 | 65 | 48 | 82 | 32 |
| Bacteroides thetaiotaomicron 7330 | 8 | 1 | 2 | 0 |
| Bacteroides thetaiotaomicron VPI-5482 | 25 | 1 | 4 | 2 |
| Bacteroides uniformis ATCC 8492 | 118 | 31 | 40 | 24 |
| Bacteroides vulgatus ATCC 8482 | 14 | 3 | 0 | 0 |
| Bifidobacterium adolescentis L2-32 | 39 | 1 | 2 | 5 |
| Bifidobacterium bifidum ATCC 29521 | 379 | 5 | 0 | 0 |
| Bifidobacterium pseudocatenulatum DSM 20438 | 288 | 75 | 8 | 2 |
| Blautia hansenii DSM 20583 | 95 | 1 | 0 | 0 |
| Blautia luti DSM 14534 | 70 | 30 | 26 | 18 |
| Candidatus Ruthia magnifica str. | 210 | 3 | 0 | 0 |
| Cereibacter sphaeroides ATCC 17029 | 284 | 36 | 0 | 1 |
| Citrobacter youngae ATCC 29220 | 278 | 37 | 18 | 1 |
| Clostridium bolteae ATCC BAA-613 | 1608 | 179 | 1 | 0 |
| Clostridium sp. M62/1 | 376 | 27 | 10 | 6 |
| Clostridium sporogenes ATCC 15579 | 8 | 0 | 0 | 0 |
| Collinsella intestinalis DSM 13280 | 62 | 56 | 39 | 2 |
| Collinsella stercoris DSM 13279 | 31 | 8 | 19 | 3 |
| Coprococcus comes ATCC 27758 | 205 | 42 | 25 | 6 |
| Dorea formicigenerans ATCC 27755 | 83 | 25 | 21 | 6 |
| Edwardsiella tarda ATCC 23685 | 424 | 30 | 12 | 3 |
| Enterobacter cancerogenus ATCC 35316 | 677 | 50 | 2 | 3 |
| Escherichia fergusonii ATCC 35469 | 0 | 3 | 1 | 0 |
| Geotalea uraniireducens Rf4 | 6 | 0 | 0 | 0 |
| Gluconacetobacter diazotrophicus PA1 5 | 13 | 0 | 1 | 0 |
| Granulicella tundricola MP5ACTX9 | 160 | 34 | 14 | 3 |
| Herpetosiphon aurantiacus DSM 785 | 1490 | 138 | 2 | 0 |
| Holdemanella biformis DSM 3989 | 262 | 50 | 33 | 1 |
| Holdemania filiformis DSM 12042 | 155 | 95 | 92 | 5 |
| Hungatella hathewayi DSM 13479 | 405 | 320 | 177 | 5 |
| Lachnospira eligens ATCC 27750 | 71 | 20 | 35 | 12 |
| Lacticaseibacillus paracasei ATCC 334 | 76 | 0 | 1 | 0 |
| Lactobacillus reuteri DSM 20016 | 3 | 0 | 0 | 0 |
| Lactococcus lactis subsp. Lactis | 15 | 8 | 3 | 0 |
| Leptothrix cholodnii SP-6 | 49 | 0 | 0 | 0 |
| Ligilactobacillus ruminis DSM 20403 = NBRC 102161 | 14 | 34 | 53 | 8 |
| Maricaulis maris MCS10 | 808 | 167 | 0 | 0 |
| Marvinbryantia formatexigens DSM 14469 | 47 | 41 | 58 | 24 |
| Megamonas funiformis YIT 11815 | 72 | 10 | 10 | 3 |
| Methylorubrum extorquens CM4 | 64 | 34 | 3 | 0 |
| Methylovorus glucosotrophus | 0 | 1 | 0 | 1 |
| Nitratifractor salsuginis DSM 16511 | 24 | 0 | 0 | 0 |
| Nocardiopsis dassonvillei subsp. dassonvillei DSM 43111 | 87 | 0 | 0 | 0 |
| Parabacteroides johnsonii DSM 18315 | 278 | 93 | 21 | 0 |
| Parabacteroides merdae ATCC 43184 | 215 | 70 | 30 | 12 |
| Pelodictyon phaeoclathratiforme BU-1 | 5 | 0 | 0 | 0 |
| Phocaeicola dorei DSM 17855 | 286 | 51 | 43 | 7 |
| Polynucleobacter necessarius subsp. necessarius STIR1 | 1 | 0 | 1 | 2 |
| Porphyromonas asaccharolytica DSM 20707 | 316 | 2 | 0 | 0 |
| Proteus penneri ATCC 35198 | 509 | 145 | 9 | 3 |
| Rhizobium leguminosarum bv. trifolii WSM1325 | 27 | 1 | 3 | 7 |
| Roseburia intestinalis L1-82 | 201 | 85 | 98 | 8 |
| Ruminiclostridium cellulolyticum H10 | 2 | 0 | 0 | 0 |
| Ruminococcus gnavus ATCC 29149 | 105 | 35 | 33 | 21 |
| Salinispora arenicola CNS-205 | 719 | 9 | 0 | 0 |
| Sediminispirochaeta smaragdinae DSM 11293 | 4 | 0 | 0 | 0 |
| Shewanella baltica OS117 | 86 | 5 | 4 | 2 |
| Solitalea canadensis DSM 3403 | 4 | 0 | 0 | 0 |
| Sphaerobacter thermophilus DSM 20745 | 224 | 106 | 7 | 0 |
| Sphaerochaeta pleomorpha str. Grapes | 882 | 163 | 1 | 0 |
| Stanieria cyanosphaera PCC 7437 | 76 | 5 | 5 | 4 |
| Staphylococcus aureus subsp. aureus JH9 | 1 | 0 | 1 | 0 |
| Streptococcus infantarius ATCC BAA-102 | 159 | 16 | 13 | 8 |
| Subdoligranulum variabile DSM 15176 | 92 | 16 | 17 | 0 |
| Syntrophobacter fumaroxidans MPOB | 3 | 0 | 0 | 0 |
| Thermoanaerobacterium thermosaccharolyticum DSM 571 | 667 | 97 | 2 | 0 |
| Thermus thermophilus SG0.5JP17-16 | 20 | 33 | 56 | 9 |
| Thomasclavelia ramosa DSM 1402 | 298 | 102 | 34 | 4 |
| Trichormus variabilis ATCC 29413 | 34 | 17 | 16 | 0 |
| Tsukamurella paurometabola DSM 20162 | 3 | 1 | 4 | 0 |
| Weeksella virosa DSM 16922 | 10 | 0 | 0 | 0 |
| Xylanimonas cellulosilytica DSM 15894 | 2 | 0 | 0 | 1 |

**Table S9** List of organisms with the number of contigs and the distribution of contigs’ length ranges of Sim92 dataset

| **Organism Name** | **Number of contigs** | | | |
| --- | --- | --- | --- | --- |
|  | **1000 - 3500**  **bps** | **3501 – 10000**  **bps** | **10001 – 50000**  **bps** | **> 50000**  **bps** |
| [Clostridium] asparagiforme DSM 15981 | 66 | 13 | 17 | 4 |
| [Clostridium] hylemonae DSM 15053 | 138 | 6 | 1 | 0 |
| [Clostridium] nexile DSM 1787 | 282 | 91 | 27 | 1 |
| [Clostridium] symbiosum ATCC 14940 | 54 | 81 | 156 | 11 |
| Acetivibrio thermocellus DSM 2360 | 36 | 0 | 2 | 0 |
| Acidobacterium capsulatum ATCC 51196 | 5 | 0 | 0 | 0 |
| Alistipes indistinctus YIT 12060 | 295 | 67 | 0 | 0 |
| Bacteroides cellulosilyticus DSM 14838 | 709 | 116 | 15 | 0 |
| Bacteroides finegoldii DSM 17565 | 265 | 125 | 63 | 15 |
| Bacteroides intestinalis DSM 17393 | 161 | 31 | 2 | 1 |
| Bacteroides ovatus strain ATCC 8483 | 52 | 0 | 0 | 0 |
| Bacteroides plebeius DSM 17135 | 279 | 42 | 24 | 6 |
| Bacteroides stercoris ATCC 43183 | 127 | 14 | 35 | 9 |
| Bacteroides thetaiotaomicron 3731 | 83 | 54 | 82 | 26 |
| Bacteroides thetaiotaomicron 7330 | 12 | 0 | 2 | 0 |
| Bacteroides thetaiotaomicron strain DSM 2079 | 7 | 0 | 0 | 0 |
| Bacteroides thetaiotaomicron VPI-5482 | 6 | 0 | 2 | 0 |
| Bacteroides uniformis ATCC 8492 | 135 | 35 | 38 | 19 |
| Bacteroides vulgatus ATCC 8482 | 52 | 3 | 0 | 0 |
| Bifidobacterium adolescentis L2-32 | 174 | 12 | 15 | 3 |
| Bifidobacterium angulatum DSM 20098 = JCM 7096 | 14 | 0 | 0 | 0 |
| Bifidobacterium bifidum ATCC 29521 | 119 | 1 | 0 | 0 |
| Bifidobacterium pseudocatenulatum DSM 20438 | 158 | 102 | 40 | 5 |
| Blautia hansenii DSM 20583 | 3 | 6 | 0 | 0 |
| Blautia luti DSM 14534 | 39 | 19 | 19 | 15 |
| Caldicellulosiruptor bescii DSM 6725 | 0 | 2 | 0 | 0 |
| Caldicellulosiruptor saccharolyticus DSM 8903 | 5 | 0 | 0 | 0 |
| Candidatus Aciduliprofundum boonei | 2 | 7 | 33 | 7 |
| Chlorobium phaeovibrioides | 96 | 43 | 26 | 8 |
| Citrobacter youngae ATCC 29220 | 91 | 102 | 36 | 8 |
| Clostridium bolteae ATCC BAA-613 | 1 | 1 | 2 | 0 |
| Clostridium sp. M62/1 | 98 | 49 | 27 | 12 |
| Collinsella intestinalis DSM 13280 | 14 | 2 | 8 | 2 |
| Collinsella stercoris DSM 13279 | 276 | 26 | 11 | 1 |
| Coprococcus comes ATCC 27758 | 138 | 98 | 32 | 10 |
| Deinococcus radiodurans R1 = ATCC 13939 = DSM 20539 strain | 82 | 3 | 7 | 0 |
| Desulfovibrio piger | 198 | 88 | 31 | 3 |
| Desulfovibrio vulgaris str. 'Miyazaki F' | 0 | 1 | 3 | 1 |
| Dictyoglomus turgidum DSM 6724 | 15 | 0 | 0 | 0 |
| Dorea formicigenerans ATCC 27755 | 83 | 15 | 16 | 10 |
| Edwardsiella tarda ATCC 23685 | 52 | 31 | 26 | 3 |
| Enterobacter cancerogenus ATCC 35316 | 1022 | 159 | 3 | 1 |
| Enterococcus faecalis EnGen0336 | 62 | 1 | 0 | 1 |
| Escherichia fergusonii ATCC 35469 | 1 | 1 | 1 | 0 |
| Haloferax volcanii DS2 | 103 | 24 | 17 | 0 |
| Herpetosiphon aurantiacus DSM 785 | 63 | 8 | 17 | 0 |
| Holdemanella biformis DSM 3989 | 159 | 17 | 36 | 3 |
| Holdemania filiformis DSM 12042 | 370 | 72 | 55 | 3 |
| Hungatella hathewayi DSM 13479 | 454 | 318 | 165 | 5 |
| Hydrogenobaculum sp. Y04AAS1 | 182 | 5 | 0 | 0 |
| Lachnospira eligens ATCC 27750 | 3 | 3 | 4 | 4 |
| Ligilactobacillus ruminis DSM 20403 = NBRC 102161 | 24 | 45 | 57 | 7 |
| Marvinbryantia formatexigens DSM 14469 | 142 | 86 | 65 | 15 |
| Megamonas funiformis YIT 11815 | 58 | 33 | 15 | 2 |
| Methanocaldococcus jannaschii DSM 2661 | 9 | 2 | 2 | 0 |
| Methanococcus maripaludis C5 | 3 | 1 | 0 | 0 |
| Methanococcus maripaludis strain S2 | 28 | 0 | 0 | 0 |
| Methanopyrus kandleri AV19 | 2 | 0 | 0 | 0 |
| Nanoarchaeum equitans Kin4-M | 18 | 19 | 18 | 0 |
| Nitrosomonas europaea ATCC 19718 | 3 | 0 | 0 | 0 |
| Nostoc sp. PCC 7120 = FACHB-418 | 249 | 89 | 93 | 30 |
| Parabacteroides johnsonii DSM 18315 | 375 | 98 | 28 | 0 |
| Parabacteroides merdae ATCC 43184 | 316 | 83 | 35 | 7 |
| Paraburkholderia xenovorans LB400 | 448 | 12 | 0 | 0 |
| Persephonella marina EX-H1 | 26 | 0 | 0 | 1 |
| Phocaeicola dorei DSM 17855 | 527 | 107 | 29 | 7 |
| Phocaeicola vulgatus MG01-10 DNA | 91 | 10 | 0 | 0 |
| Porphyromonas gingivalis ATCC 33277 | 263 | 1 | 0 | 0 |
| Proteus penneri ATCC 35198 | 207 | 38 | 9 | 0 |
| Pyrobaculum aerophilum str. IM2 | 64 | 1 | 0 | 0 |
| Pyrobaculum arsenaticum DSM 13514 | 460 | 12 | 0 | 0 |
| Pyrobaculum calidifontis JCM 11548 | 11 | 0 | 0 | 0 |
| Pyrococcus furiosus DSM 3638 | 40 | 0 | 0 | 0 |
| Roseburia intestinalis L1-82 | 221 | 114 | 111 | 8 |
| Ruegeria pomeroyi DSS-3 | 1 | 7 | 14 | 2 |
| Ruminococcus gnavus ATCC 29149 | 162 | 65 | 31 | 19 |
| Shewanella baltica OS185 | 415 | 7 | 1 | 0 |
| Shewanella baltica OS223 | 26 | 11 | 2 | 0 |
| Streptococcus infantarius ATCC BAA-102 | 51 | 11 | 10 | 11 |
| Subdoligranulum variabile DSM 15176 | 208 | 124 | 83 | 3 |
| Sulfitobacter sp. EE-36 | 19 | 5 | 2 | 2 |
| Sulfitobacter sp. NAS-14.1 | 50 | 22 | 6 | 4 |
| Sulfolobus tokodaii str. 7 DNA | 5 | 0 | 0 | 0 |
| Sulfurihydrogenibium yellowstonense SS-5 | 62 | 120 | 46 | 0 |
| Thermoanaerobacter pseudethanolicus ATCC 33223 | 5 | 0 | 0 | 0 |
| Thermotoga neapolitana DSM 4359 | 205 | 2 | 0 | 0 |
| Thermotoga petrophila RKU-1 | 41 | 1 | 0 | 0 |
| Thermotoga sp. RQ2 | 43 | 46 | 61 | 4 |
| Thermus thermophilus HB8 | 37 | 2 | 5 | 2 |
| Thomasclavelia ramosa DSM 1402 | 369 | 104 | 27 | 3 |
| Treponema denticola ATCC 35405 | 142 | 169 | 89 | 0 |
| Wolinella succinogenes DSM 1740 | 171 | 1 | 0 | 0 |

**Table S10** List of organisms with the number of contigs and the distribution of contigs’ length ranges of Sim98 dataset

| **Organism Name** | **Number of contigs** | | | |
| --- | --- | --- | --- | --- |
|  | **1000 - 3500**  **bps** | **3501 – 10000**  **bps** | **10001 – 50000**  **bps** | **> 50000**  **bps** |
| [Clostridium] asparagiforme DSM 15981 | 798 | 57 | 5 | 1 |
| [Clostridium] hylemonae DSM 15053 | 199 | 37 | 1 | 0 |
| [Clostridium] nexile DSM 1787 | 281 | 87 | 38 | 5 |
| [Clostridium] symbiosum ATCC 14940 | 87 | 102 | 162 | 5 |
| Akkermansia muciniphila | 1 | 0 | 2 | 0 |
| Akkermansia muciniphila ATCC BAA-835 | 6 | 0 | 0 | 0 |
| Alistipes finegoldii DSM 17242 | 341 | 6 | 0 | 0 |
| Alistipes indistinctus YIT 12060 | 17 | 0 | 0 | 0 |
| Anaerobutyricum hallii DSM 3353 | 52 | 67 | 102 | 6 |
| Anaerostipes hadrus ATCC 29173 = JCM 17467 | 0 | 0 | 2 | 0 |
| Bacteroides cellulosilyticus DSM 14838 | 752 | 97 | 31 | 0 |
| Bacteroides finegoldii DSM 17565 | 179 | 107 | 80 | 11 |
| Bacteroides intestinalis DSM 17393 | 274 | 54 | 1 | 0 |
| Bacteroides ovatus strain ATCC 8483 | 4 | 1 | 0 | 0 |
| Bacteroides plebeius DSM 17135 | 217 | 43 | 21 | 9 |
| Bacteroides stercoris ATCC 43183 | 251 | 18 | 26 | 2 |
| Bacteroides thetaiotaomicron 3731 | 233 | 77 | 68 | 29 |
| Bacteroides thetaiotaomicron 7330 | 18 | 1 | 0 | 2 |
| Bacteroides thetaiotaomicron strain DSM 2079 | 13 | 0 | 3 | 1 |
| Bacteroides thetaiotaomicron VPI-5482 | 32 | 1 | 1 | 0 |
| Bacteroides uniformis ATCC 8492 | 310 | 56 | 33 | 12 |
| Bacteroides vulgatus ATCC 8482 | 24 | 1 | 0 | 0 |
| Bifidobacterium adolescentis L2-32 | 298 | 63 | 13 | 2 |
| Bifidobacterium longum subsp. longum JCM 1217 | 167 | 33 | 20 | 6 |
| Bifidobacterium pseudocatenulatum DSM 20438 | 167 | 54 | 11 | 2 |
| Blautia hansenii DSM 20583 | 4 | 1 | 0 | 0 |
| Blautia luti DSM 14534 | 205 | 95 | 69 | 5 |
| Blautia wexlerae DSM 19850 | 143 | 93 | 111 | 10 |
| Caldicellulosiruptor bescii DSM 6725 | 3 | 2 | 0 | 0 |
| Caldicellulosiruptor saccharolyticus DSM 8903 | 6 | 0 | 0 | 0 |
| Candidatus Aciduliprofundum boonei | 1 | 7 | 28 | 8 |
| Chlorobium phaeobacteroides DSM 266 | 259 | 4 | 0 | 0 |
| Chlorobium phaeovibrioides | 25 | 13 | 19 | 12 |
| Chlorobium tepidum TLS | 305 | 4 | 0 | 0 |
| Chloroflexus aurantiacus J-10-fl | 6 | 0 | 0 | 0 |
| Citrobacter youngae ATCC 29220 | 79 | 9 | 7 | 0 |
| Clostridium bolteae ATCC BAA-613 | 3 | 0 | 1 | 0 |
| Clostridium butyricum DSM 10702 | 165 | 52 | 50 | 25 |
| Clostridium sp. M62/1 | 359 | 66 | 21 | 4 |
| Collinsella aerofaciens | 62 | 26 | 35 | 9 |
| Collinsella intestinalis DSM 13280 | 12 | 21 | 10 | 0 |
| Collinsella stercoris DSM 13279 | 320 | 49 | 15 | 3 |
| Coprococcus comes ATCC 27758 | 322 | 53 | 13 | 3 |
| Deinococcus radiodurans R1 = ATCC 13939 = DSM 20539 strain | 71 | 2 | 6 | 1 |
| Desulfovibrio piger | 358 | 125 | 24 | 4 |
| Dictyoglomus turgidum DSM 6724 | 137 | 1 | 0 | 0 |
| Dorea formicigenerans ATCC 27755 | 81 | 69 | 32 | 8 |
| Edwardsiella tarda ATCC 23685 | 44 | 28 | 22 | 3 |
| Enterobacter cancerogenus ATCC 35316 | 0 | 2 | 4 | 3 |
| Enterococcus faecalis EnGen0336 | 106 | 1 | 0 | 1 |
| Escherichia fergusonii ATCC 35469 | 0 | 1 | 2 | 0 |
| Fusobacterium nucleatum subsp. | 3 | 0 | 1 | 0 |
| Haloferax volcanii DS2 | 59 | 38 | 19 | 0 |
| Herpetosiphon aurantiacus DSM 785 | 3 | 3 | 12 | 1 |
| Holdemanella biformis DSM 3989 | 79 | 53 | 64 | 5 |
| Holdemania filiformis DSM 12042 | 190 | 61 | 76 | 4 |
| Hungatella hathewayi DSM 13479 | 461 | 337 | 159 | 4 |
| Hydrogenobaculum sp. Y04AAS1 | 8 | 0 | 0 | 0 |
| Lachnospira eligens ATCC 27750 | 3 | 3 | 4 | 4 |
| Lactobacillus gasseri ATCC 33323 = JCM 1131 | 48 | 1 | 0 | 0 |
| Lactobacillus reuteri DSM 20016 | 6 | 0 | 0 | 0 |
| Ligilactobacillus ruminis DSM 20403 = NBRC 102161 | 52 | 52 | 57 | 5 |
| Marvinbryantia formatexigens DSM 14469 | 261 | 69 | 54 | 15 |
| Megamonas funiformis YIT 11815 | 286 | 8 | 13 | 1 |
| Methanocaldococcus jannaschii DSM 2661 | 20 | 2 | 2 | 0 |
| Methanococcus maripaludis strain S2 | 270 | 9 | 0 | 0 |
| Nanoarchaeum equitans Kin4-M | 0 | 1 | 10 | 3 |
| Nitrosomonas europaea ATCC 19718 | 3 | 0 | 0 | 0 |
| Nostoc sp. PCC 7120 = FACHB-418 | 406 | 103 | 65 | 34 |
| Parabacteroides johnsonii DSM 18315 | 420 | 88 | 17 | 0 |
| Parabacteroides merdae ATCC 43184 | 243 | 125 | 48 | 2 |
| Paraburkholderia xenovorans LB400 | 288 | 7 | 0 | 0 |
| Phocaeicola dorei DSM 17855 | 143 | 90 | 32 | 8 |
| Phocaeicola vulgatus MG01-10 DNA | 107 | 4 | 0 | 0 |
| Prevotella melaninogenica ATCC 25845 | 184 | 2 | 0 | 0 |
| Proteus penneri ATCC 35198 | 332 | 40 | 18 | 8 |
| Pyrococcus horikoshii OT3 | 131 | 0 | 0 | 0 |
| Roseburia intestinalis L1-82 | 208 | 112 | 87 | 5 |
| Ruegeria pomeroyi DSS-3 | 122 | 8 | 0 | 0 |
| Ruminococcus gauvreauii DSM 19829 | 187 | 69 | 57 | 15 |
| Ruminococcus gnavus ATCC 29149 | 70 | 36 | 36 | 18 |
| Salinispora tropica CNB-440 | 2 | 0 | 0 | 0 |
| Segatella copri DSM 18205 | 167 | 89 | 38 | 8 |
| Shewanella baltica OS185 | 7 | 3 | 0 | 0 |
| Shewanella baltica OS223 | 10 | 7 | 2 | 0 |
| Streptococcus infantarius ATCC BAA-102 | 210 | 36 | 15 | 5 |
| Subdoligranulum variabile DSM 15176 | 80 | 8 | 12 | 1 |
| Sulfitobacter sp. EE-36 | 53 | 6 | 3 | 0 |
| Sulfitobacter sp. NAS-14.1 | 40 | 20 | 22 | 3 |
| Sulfurihydrogenibium sp. YO3AOP1 | 71 | 8 | 0 | 0 |
| Sulfurihydrogenibium yellowstonense SS-5 | 38 | 93 | 48 | 0 |
| Thermoanaerobacter pseudethanolicus ATCC 33223 | 9 | 1 | 0 | 0 |
| Thermotoga neapolitana DSM 4359 | 16 | 0 | 0 | 0 |
| Thermotoga petrophila RKU-1 | 6 | 0 | 0 | 0 |
| Thermotoga sp. RQ2 | 17 | 0 | 0 | 0 |
| Thermus thermophilus HB8 | 2 | 2 | 5 | 2 |
| Thomasclavelia ramosa DSM 1402 | 201 | 79 | 42 | 4 |
| Wolinella succinogenes DSM 1740 | 2 | 0 | 0 | 0 |

**Table S11** List of organisms with the number of contigs and the distribution of contigs’ length ranges of SRR8073716 dataset

| **Organism Name** | **Number of contigs** | | | |
| --- | --- | --- | --- | --- |
|  | **1000 - 3500**  **bps** | **3501 – 10000**  **bps** | **10001 – 50000 bps** | **> 50000**  **bps** |
| Cohaesibacter sp. ES.047 | 31 | 22 | 40 | 26 |
| Halomonas sp. HL-4 | 62 | 40 | 22 | 1 |
| Halomonas sp. HL-93 | 75 | 51 | 35 | 0 |
| Marinobacter sp. LV10MA510-1 | 39 | 21 | 32 | 22 |
| Marinobacter sp. LV10R510-8 | 33 | 14 | 13 | 30 |
| Micromonospora coxensis DSM 45161 | 2 | 0 | 0 | 0 |
| Micromonospora echinaurantiaca DSM 43904 | 12 | 5 | 15 | 29 |
| Micromonospora echinofusca DSM 43913 | 14 | 4 | 11 | 35 |
| Muricauda sp. ES.050 | 4 | 2 | 3 | 7 |
| Propionibacteriaceae bacterium ES.041 | 1 | 0 | 1 | 7 |
| Psychrobacter sp. LV10R520-6 | 20 | 10 | 33 | 22 |
| Thioclava sp. ES.032 | 11 | 4 | 20 | 22 |

**Table S12** List of organisms with the number of contigs and the distribution of contigs’ length ranges of SRR11487931 dataset

| **Organism Name** | **Number of contigs** | | | |
| --- | --- | --- | --- | --- |
|  | **1000 - 3500**  **bps** | **3501 – 10000**  **bps** | **10001 – 50000 bps** | **> 50000**  **bps** |
| Akkermansia muciniphila | 28 | 0 | 0 | 0 |
| Alistipes finegoldii | 1109 | 89 | 0 | 0 |
| Eubacterium hallii | 457 | 4 | 0 | 0 |
| Anaerostipes hadrus | 6 | 1 | 0 | 0 |
| Bacteroides thetaiotaomicron | 170 | 162 | 202 | 7 |
| Bacteroides uniformis | 35 | 0 | 0 | 0 |
| Bifidobacterium longum subsp. infantis | 84 | 41 | 26 | 0 |
| Bifidobacterium longum subsp. longum | 27 | 12 | 23 | 20 |
| Blautia wexlerae | 14 | 1 | 0 | 0 |
| Clostridium butyricum | 870 | 53 | 1 | 0 |
| Collinsella aerofaciens | 481 | 184 | 8 | 0 |
| Escherichia coli | 9 | 2 | 4 | 0 |
| Faecalibacterium prausnitzii | 12 | 1 | 0 | 0 |
| Lactobacillus gasseri | 7 | 0 | 0 | 0 |
| Parabacteroides distasonis | 33 | 40 | 88 | 30 |
| Prevotella copri | 47 | 76 | 105 | 6 |
| Prevotella melaninogenica | 597 | 210 | 14 | 0 |
| Roseburia hominis | 949 | 47 | 0 | 0 |
| Roseburia intestinalis | 11 | 0 | 0 | 0 |
| Ruminococcus gauvreauii | 3 | 0 | 0 | 0 |

**Table S13** List of organisms with the number of contigs and the distribution of contigs’ length ranges of SRR11487935 dataset

| **Organism Name** | **Number of contigs** | | | |
| --- | --- | --- | --- | --- |
|  | **1000 - 3500**  **bps** | **3501 – 10000**  **bps** | **10001 – 50000 bps** | **> 50000**  **bps** |
| Akkermansia muciniphila | 17 | 143 | 0 | 0 |
| Alistipes finegoldii | 1036 | 0 | 4 | 0 |
| Eubacterium hallii | 7 | 1 | 0 | 0 |
| Anaerostipes hadrus | 165 | 191 | 199 | 4 |
| Bacteroides thetaiotaomicron | 53 | 0 | 1 | 0 |
| Bacteroides uniformis | 84 | 42 | 25 | 0 |
| Bifidobacterium longum subsp. infantis | 27 | 13 | 16 | 23 |
| Bifidobacterium longum subsp. longum | 12 | 0 | 0 | 0 |
| Blautia wexlerae | 885 | 34 | 2 | 0 |
| Clostridium butyricum | 405 | 205 | 10 | 0 |
| Collinsella aerofaciens | 10 | 5 | 5 | 0 |
| Escherichia coli | 467 | 6 | 0 | 0 |
| Faecalibacterium prausnitzii | 16 | 0 | 0 | 0 |
| Lactobacillus gasseri | 7 | 0 | 0 | 0 |
| Parabacteroides distasonis | 37 | 34 | 77 | 32 |
| Prevotella copri | 58 | 85 | 100 | 7 |
| Prevotella melaninogenica | 562 | 210 | 16 | 0 |
| Roseburia hominis | 946 | 64 | 0 | 0 |
| Roseburia intestinalis | 10 | 0 | 0 | 0 |
| Ruminococcus gauvreauii | 15 | 0 | 0 | 0 |

**Table S14** List of organisms with the number of contigs and the distribution of contigs’ length ranges of SRR3656745 dataset

| **Organism Name** | **Number of contigs** | | | |
| --- | --- | --- | --- | --- |
|  | **1000 - 3500**  **bps** | **3501 – 10000**  **bps** | **10001 – 50000 bps** | **> 50000**  **bps** |
| Clostridium perfringens ATCC 13124 | 2 | 3 | 2 | 10 |
| Clostridium thermocellum ATCC 27405 | 32 | 16 | 38 | 25 |
| Coraliomargarita akajimensis DSM 45221 | 5 | 7 | 2 | 9 |
| Corynebacterium glutamicum ATCC 13032 | 11 | 3 | 11 | 19 |
| Desulfosporosinus acidiphilus SJ4 DSM 22704 | 17 | 10 | 25 | 31 |
| Desulfosporosinus meridiei DSM 13257 | 7 | 5 | 7 | 28 |
| Desulfotomaculum gibsoniae DSM 7213 | 23 | 9 | 35 | 29 |
| Escherichia coli K-12, MG1655 | 27 | 18 | 46 | 33 |
| Echinicola vietnamensis DSM 17526 | 16 | 3 | 17 | 32 |
| Fervidobacterium pennivorans DSM 9078 | 8 | 9 | 11 | 17 |
| Frateuria aurantia DSM 6220 | 2 | 1 | 0 | 6 |
| Halovivax ruber XH-70 | 1 | 0 | 3 | 5 |
| Hirschia baltica ATCC 49814 | 1 | 1 | 0 | 4 |
| Meiothermus Silvanus DSM 9946 | 59 | 49 | 105 | 10 |
| Natronobacterium gregoryi SP2 | 19 | 18 | 56 | 21 |
| Natronococcus occultus DSM 3396 | 0 | 0 | 3 | 11 |
| Nocardiopsis dassonvillei DSM 43111 | 2 | 0 | 0 | 0 |
| Olsenella uli DSM 7084 | 0 | 1 | 1 | 3 |
| Pseudomonas stutzeri RCH2 | 5 | 4 | 3 | 10 |
| Salmonella bongori NCTC 12419 | 95 | 106 | 89 | 18 |
| Salmonella enterica subsp. Arizonae serovar RSK2980 | 32 | 13 | 38 | 35 |
| Segniliparus rotundus DSM 44985 | 6 | 0 | 2 | 10 |
| Spirochaeta smaragdinae DSM 11293 | 6 | 1 | 2 | 17 |
| Streptococcus pyogenes M1 GAS SF370 | 6 | 0 | 7 | 11 |
| Terriglobus roseus DSM 18391 | 2 | 1 | 3 | 3 |
| Thermobacillus composti KWC4, DSM 18247 | 41 | 20 | 44 | 31 |

**Table S15** List of organisms with the number of contigs and the distribution of contigs’ length ranges of SRR606249 dataset

| **Organism** | **Number of contigs** | | | |
| --- | --- | --- | --- | --- |
|  | **1000 - 3500**  **bps** | **3501 – 10000**  **bps** | **10001 – 50000 bps** | **> 50000**  **bps** |
| Acidobacterium capsulatum | 4 | 5 | 4 | 19 |
| Aciduliprofundum boonei | 2 | 4 | 9 | 6 |
| Akkermansia muciniphila | 2 | 9 | 2 | 0 |
| Archaeoglobus fulgidus | 5 | 0 | 5 | 2 |
| Bacteroides thetaiotaomicron | 22 | 13 | 23 | 37 |
| Bacteroides vulgatus | 9 | 13 | 12 | 1 |
| Bordetella bronchiseptica | 605 | 348 | 78 | 0 |
| Burkholderia xenovorans LB400 | 1366 | 628 | 140 | 2 |
| Caldicellulosiruptor bescii | 60 | 37 | 46 | 16 |
| Caldicellulosiruptor saccharolyticus | 74 | 48 | 46 | 16 |
| Chlorobium limicola | 20 | 14 | 17 | 19 |
| Chlorobium phaeobacteroides | 39 | 28 | 56 | 14 |
| Chlorobium phaeovibrioides | 0 | 2 | 2 | 0 |
| Chlorobium tepidum | 1 | 2 | 7 | 11 |
| Chloroflexus aurantiacus J-10-fl | 22 | 24 | 65 | 31 |
| Clostridium thermocellum | 16 | 11 | 25 | 7 |
| Deinococcus radiodurans R1 | 103 | 95 | 87 | 2 |
| Desulfovibrio piger | 4 | 6 | 16 | 11 |
| Desulfovibrio vulgaris DP4 | 3 | 5 | 4 | 0 |
| Dictyoglomus turgidum | 5 | 0 | 1 | 7 |
| Enterococcus faecalis | 3 | 0 | 0 | 0 |
| Fusobacterium nucleatum | 5 | 6 | 2 | 0 |
| Gemmatimonas aurantiaca | 2 | 1 | 2 | 8 |
| Geobacter sulfurreducens PCA | 17 | 3 | 8 | 19 |
| Haloferax volcanii | 50 | 77 | 121 | 10 |
| Herpetosiphon aurantiacus | 43 | 20 | 40 | 41 |
| Hydrogenobaculum sp. Y04AAS1 | 3 | 2 | 1 | 11 |
| Ignicoccus hospitalis | 1 | 2 | 5 | 7 |
| Leptothrix cholodnii | 196 | 185 | 152 | 5 |
| Methanocaldococcus jannaschii | 3 | 5 | 12 | 7 |
| Methanococcus maripaludis C5 | 8 | 6 | 10 | 9 |
| Methanococcus maripaludis S2 | 3 | 1 | 7 | 11 |
| Methanopyrus kandleri | 0 | 1 | 0 | 2 |
| Methanosarcina acetivorans C2A | 90 | 116 | 149 | 14 |
| Nanoarchaeum equitans | 0 | 0 | 0 | 1 |
| Nitrosomonas europaea | 21 | 15 | 48 | 13 |
| Nostoc sp. PCC 7120 | 20 | 19 | 37 | 43 |
| Pelodictyon phaeoclathratiforme | 19 | 16 | 19 | 23 |
| Persephonella marina EX-H1 | 0 | 1 | 0 | 4 |
| Porphyromonas gingivalis | 16 | 24 | 56 | 9 |
| Pyrobaculum aerophilum IM2 | 8 | 11 | 20 | 9 |
| Pyrobaculum arsenaticum | 2 | 3 | 7 | 13 |
| Pyrobaculum calidifontis | 1 | 1 | 3 | 8 |
| Pyrococcus furiosus | 5 | 8 | 21 | 17 |
| Pyrococcus horikoshii | 1 | 2 | 3 | 6 |
| Rhodopirellula baltica | 19 | 9 | 18 | 41 |
| Ruegeria pomeroyi | 210 | 188 | 128 | 5 |
| Salinispora arenicola | 254 | 207 | 48 | 0 |
| Salinispora tropica | 343 | 267 | 129 | 6 |
| Shewanella baltica OS185 | 816 | 346 | 39 | 0 |
| Shewanella baltica OS223 | 360 | 68 | 6 | 0 |
| Sulfitobacter sp. EE-36 | 482 | 395 | 81 | 3 |
| Sulfitobacter sp. NAS-14.1 | 2 | 0 | 0 | 0 |
| Sulfolobus tokodaii | 14 | 12 | 28 | 14 |
| Sulfurihydrogenibium sp. YO3AOP1 | 143 | 57 | 37 | 5 |
| Sulfurihydrogenibium yellowstonense SS-5 | 70 | 16 | 1 | 0 |
| Thermoanaerobacter pseudethanolicus | 19 | 12 | 30 | 12 |
| Thermotoga neapolitana DSM 4359 | 13 | 17 | 25 | 9 |
| Thermotoga petrophila RKU-1 | 45 | 15 | 7 | 0 |
| Thermotoga sp. RQ2 | 39 | 26 | 40 | 10 |
| Thermus thermophilus HB8 | 11 | 14 | 29 | 13 |
| Treponema denticola | 18 | 6 | 6 | 16 |
| Wolinella succinogenes | 7 | 6 | 11 | 12 |
| Zymomonas mobilis | 15 | 22 | 37 | 13 |

**Table S16** Clustering results of 1000 to 1000000 bps contigs, with varying coverage and without partitioning contigs into groups.

|  | **Dataset** | **Ground**  **truth**  **bins** | **Number of bins identified** | | |
| --- | --- | --- | --- | --- | --- |
|  |  |  | **1.4 -fold coverage difference** | **1.5 -fold**  **coverage difference** | **1.6 -fold**  **coverage difference** |
| Simulated datasets | Sim-5G | 5 | 10 | **9** | **9** |
|  | SRR8304764 | 51 | **52** | 42 | 41 |
|  | SRR8304773 | 51 | 57 | **46** | 40 |
|  | SRR8304775 | 51 | **43** | 34 | 33 |
|  | SRR8304776 | 51 | **52** | 40 | 32 |
|  | Sim79 | 79 | **82** | 61 | 58 |
|  | Sim82 | 82 | **83** | 78 | 75 |
|  | Sim92 | 92 | 104 | **87** | 78 |
|  | Sim98 | 98 | 112 | **93** | 75 |
| Mock community datasets | SRR8073716 | 12 | 16 | **15** | **15** |
|  | SRR11487931 | 20 | **21** | **19** | **19** |
|  | SRR11487935 | 20 | **21** | **21** | 18 |
|  | SRR3656745 | 26 | 28 | **25** | **25** |
|  | SRR606249 | 64 | **62** | 53 | 52 |

The best result in terms of the number of bins identified is highlighted in bold red.

**Table S17** Clustering results based on partitioning contigs into two groups according to their lengths and varying coverage information:

Group 1: consists of contigs ranging from 1000 to 9000 bps.

Group 2: consists of contigs ranging from 6500 to 1000000 bps.

|  | **Dataset** | **Ground**  **truth**  **bins** | **Number of bins identified** | | |
| --- | --- | --- | --- | --- | --- |
|  |  |  | **1.4 -fold coverage difference** | **1.5 -fold**  **coverage difference** | **1.6 -fold**  **coverage difference** |
| Simulated datasets | Sim-5G | 5 | **11** | **11** | **11** |
|  | SRR8304764 | 51 | 59 | **50** | 40 |
|  | SRR8304773 | 51 | 64 | **53** | 47 |
|  | SRR8304775 | 51 | **47** | 41 | 38 |
|  | SRR8304776 | 51 | **52** | 43 | 37 |
|  | Sim79 | 79 | 99 | **82** | 65 |
|  | Sim82 | 82 | 101 | **85** | 78 |
|  | Sim92 | 92 | 112 | 103 | **90** |
|  | Sim98 | 98 | 138 | 108 | **95** |
| Mock community datasets | SRR8073716 | 12 | 25 | 25 | **22** |
|  | SRR11487931 | 20 | 22 | **19** | **19** |
|  | SRR11487935 | 20 | 26 | **22** | **22** |
|  | SRR3656745 | 26 | 34 | 31 | **29** |
|  | SRR606249 | 64 | **58** | 53 | 48 |

The best result in terms of the number of bins identified is highlighted in bold red.

**Table S18** Clustering results based on partitioning contigs into three groups according to their lengths and varying coverage information:

Group 1: consists of contigs ranging from 1000 to 9000 bps.

Group 2: consists of contigs ranging from 3500 to 25000 bps.

Group 3: consists of contigs ranging from 9000 to 1000000 bps.

|  | **Dataset** | **Ground**  **truth**  **bins** | **Number of bins identified** | | |
| --- | --- | --- | --- | --- | --- |
|  |  |  | **1.4 -fold coverage difference** | **1.5 -fold**  **coverage difference** | **1.6 -fold**  **coverage difference** |
| Simulated datasets | Sim-5G | 5 | **10** | **10** | **10** |
|  | SRR8304764 | 51 | 44 | **51** | 38 |
|  | SRR8304773 | 51 | 60 | **55** | 48 |
|  | SRR8304775 | 51 | 44 | **47** | 35 |
|  | SRR8304776 | 51 | 57 | **48** | 43 |
|  | Sim79 | 79 | 90 | **82** | 58 |
|  | Sim82 | 82 | 93 | **86** | 75 |
|  | Sim92 | 92 | 113 | **96** | 86 |
|  | Sim98 | 98 | 124 | **107** | 87 |
| Mock community datasets | SRR8073716 | 12 | **20** | **20** | **20** |
|  | SRR11487931 | 20 | 23 | **22** | **22** |
|  | SRR11487935 | 20 | 25 | **18** | **18** |
|  | SRR3656745 | 26 | 29 | **24** | 22 |
|  | SRR606249 | 64 | **59** | **59** | 53 |

The best result in terms of the number of bins identified is highlighted in bold red.

**Table S19** Clustering results based on partitioning contigs into four groups according to their lengths and varying coverage information:

Group 1: consists of contigs ranging from 1000 to 9000 bps.

Group 2: consists of contigs ranging from 3500 to 25000 bps.

Group 3: consists of contigs ranging from 9000 to 100000 bps.

Group 4: consists of contigs ranging from 15000 to 1000000 bps.

|  | **Dataset** | **Ground**  **truth**  **bins** | **Number of bins identified** | | |
| --- | --- | --- | --- | --- | --- |
|  |  |  | **1.4 -fold coverage difference** | **1.5 -fold**  **coverage difference** | **1.6 -fold**  **coverage difference** |
| Simulated datasets | Sim-5G | 5 | **10** | **10** | **10** |
|  | SRR8304764 | 51 | 44 | **51** | 38 |
|  | SRR8304773 | 51 | 60 | **55** | 48 |
|  | SRR8304775 | 51 | 44 | **47** | 35 |
|  | SRR8304776 | 51 | 57 | **48** | 43 |
|  | Sim79 | 79 | 90 | **82** | 58 |
|  | Sim82 | 82 | 93 | **86** | 75 |
|  | Sim92 | 92 | 113 | **96** | 86 |
|  | Sim98 | 98 | 124 | **107** | 87 |
| Mock community datasets | SRR8073716 | 12 | **20** | **20** | **20** |
|  | SRR11487931 | 20 | 23 | **22** | **22** |
|  | SRR11487935 | 20 | 25 | **18** | **18** |
|  | SRR3656745 | 26 | 29 | **24** | 22 |
|  | SRR606249 | 64 | **59** | **59** | 53 |

The best result in terms of the number of bins identified is highlighted in bold red.

**Table S20** List of organisms in Sim79 dataset

| **Organisms** | **Accession** |
| --- | --- |
| [Clostridium] asparagiforme DSM 15981 | GCA_000158075.1 |
| [Clostridium] hylemonae DSM 15053 | GCA_000156515.1 |
| [Clostridium] nexile DSM 1787 | GCA_000156035.2 |
| [Clostridium] symbiosum ATCC 14940 | GCA_000466485.1 |
| Acidithiobacillus ferrivorans SS3 | CP002985.1 |
| Acidovorax citrulli AAC00-1 | CP000512.1 |
| Actinobacillus succinogenes 130Z | CP000746.1 |
| Akkermansia muciniphila ATCC BAA-835 | GCA_017504145.1 |
| Alistipes indistinctus YIT 12060 | JH370372.1 |
| Aminobacterium colombiense DSM 12261 | CP001997.1 |
| Arcobacter nitrofigilis DSM 7299 | CP001999.1 |
| Bacillus cellulosilyticus DSM 2522 | CP002394.1 |
| Bacillus cytotoxicus NVH 391-98 | GCA_000017425.1 |
| Bacteroides cellulosilyticus DSM 14838 | GCA_000158035.1 |
| Bacteroides finegoldii DSM 17565 | GCA_000156195.1 |
| Bacteroides intestinalis DSM 17393 | GCA_000172175.1 |
| Bacteroides ovatus strain ATCC 8483 | CP012938.1 |
| Bacteroides plebeius DSM 17135 | GCA_000187895.1 |
| Bacteroides stercoris ATCC 43183 | GCA_000154525.1 |
| Bacteroides thetaiotaomicron 3731 | GCA_014131755.1 |
| Bacteroides thetaiotaomicron 7330 | CP012937.1 |
| Bacteroides thetaiotaomicron VPI-5482 | GCA_000011065.1 |
| Bacteroides uniformis ATCC 8492 | GCA_000154205.1 |
| Bacteroides vulgatus ATCC 8482 | CP000139.1 |
| Beijerinckia indica subsp. indica ATCC 9039 | GCA_000019845.1 |
| Bifidobacterium adolescentis L2-32 | GCA_000154085.1 |
| Bifidobacterium angulatum DSM 20098 | AP012322.1 |
| Bifidobacterium bifidum ATCC 29521 | AP012323.1 |
| Bifidobacterium dentium ATCC 27678 | GCA_000172135.1 |
| Bifidobacterium pseudocatenulatum DSM 20438 | GCA_000173435.1 |
| Blautia hansenii DSM 20583 | CP022413.2 |
| Blautia luti DSM 14534 | GCA_009707925.1 |
| Brevundimonas subvibrioides ATCC 15264 | CP002102.1 |
| Burkholderia orbicola MC0-3 | GCA_000019505.1 |
| Calothrix sp. PCC 6303 | GCA_000317435.1 |
| Caulobacter segnis ATCC 21756 | CP002008.1 |
| Chlamydia trachomatis G/9301 | CP001930.1 |
| Chlorobium phaeobacteroides BS1 | CP001101.1 |
| Chromohalobacter salexigens DSM 3043 | CP000285.1 |
| Citrobacter youngae ATCC 29220 | GCA_000155975.1 |
| Clostridium bolteae ATCC BAA-613 | CP022464.2/CP022465.2 |
| Clostridium lentocellum DSM 5427 | CP002582.1 |
| Clostridium sp. M62/1 | GCA_000159055.1 |
| Clostridium sporogenes ATCC 15579 | GCA_000155085.1 |
| Collinsella intestinalis DSM 13280 | GCA_000156175.1 |
| Collinsella stercoris DSM 13279 | GCA_000156215.1 |
| Coprococcus comes ATCC 27758 | GCA_000155875.1 |
| Cryptobacterium curtum DSM 15641 | CP001682.1 |
| Cyanobacterium aponinum PCC 10605 | GCA_000317675.1 |
| Cyanobium gracile PCC 6307 | CP003495.1 |
| Dehalococcoides mccartyi BAV1 | CP000688.1 |
| Desulfosporosinus acidiphilus SJ4 | GCA_000255115.3 |
| Desulfotomaculum acetoxidans DSM 771 | CP001720.1 |
| Desulfovibrio vulgaris str. 'Miyazaki F' | CP001197.1 |
| Desulfurispirillum indicum S5 | CP002432.1 |
| Dorea formicigenerans ATCC 27755 | GCA_000169235.1 |
| Edwardsiella tarda ATCC 23685 | GCA_000163955.1 |
| Eggerthella lenta DSM 2243 | CP001726.1 |
| Enterobacter cancerogenus ATCC 35316 | GCA_000155995.1 |
| Escherichia fergusonii ATCC 35469 | GCA_000026225.1 |
| Fervidobacterium nodosum Rt17-B1 | CP000771.1 |
| Fluviicola taffensis DSM 16823 | CP002542.1 |
| Holdemanella biformis DSM 3989 | GCA_000156655.1 |
| Holdemania filiformis DSM 12042 | GCA_000157995.1 |
| Hungatella hathewayi DSM 13479 | GCA_000160095.1 |
| Lachnospira eligens ATCC 27750 | GCA_000146185.1 |
| Lactobacillus reuteri DSM 20016 | CP000705.1 |
| Ligilactobacillus ruminis DSM 20403 = NBRC 102161 | GCA_001436475.1 |
| Marvinbryantia formatexigens DSM 14469 | GCA_000173815.1 |
| Megamonas funiformis YIT 11815 | GCA_000245775.1 |
| Parabacteroides johnsonii DSM 18315 | GCA_000156495.1 |
| Parabacteroides merdae ATCC 43184 | GCA_000154105.1 |
| Phocaeicola dorei DSM 17855 | GCA_000156075.1 |
| Proteus penneri ATCC 35198 | GCA_000155835.1 |
| Roseburia intestinalis L1-82 | GCA_000156535.1 |
| Ruminococcus gnavus ATCC 29149 | GCA_000169475.1 |
| Streptococcus infantarius ATCC BAA-102 | GCA_000154985.1 |
| Subdoligranulum variabile DSM 15176 | GCA_000157955.1 |
| Thomasclavelia ramosa DSM 1402 | GCA_000154485.1 |

**Table S21** List of organisms in Sim82 dataset

| **Organisms** | **Accession** |
| --- | --- |
| [Clostridium] asparagiforme DSM 15981 | GCA_000158075.1 |
| [Clostridium] hylemonae DSM 15053 | GCA_000156515.1 |
| [Clostridium] nexile DSM 1787 | GCA_000156035.2 |
| [Clostridium] symbiosum ATCC 14940 | GCA_000466485.1 |
| Alistipes indistinctus YIT 12060 | JH370372.1 |
| Bacteroides cellulosilyticus DSM 14838 | GCA_000158035.1 |
| Bacteroides finegoldii DSM 17565 | GCA_000156195.1 |
| Bacteroides intestinalis DSM 17393 | GCA_000172175.1 |
| Bacteroides plebeius DSM 17135 | GCA_000187895.1 |
| Bacteroides stercoris ATCC 43183 | GCA_000154525.1 |
| Bacteroides thetaiotaomicron 3731 | GCA_014131755.1 |
| Bacteroides thetaiotaomicron 7330 | CP012937.1 |
| Bacteroides thetaiotaomicron VPI-5482 | GCA_000011065.1 |
| Bacteroides uniformis ATCC 8492 | GCA_000154205.1 |
| Bacteroides vulgatus ATCC 8482 | CP000139.1 |
| Bifidobacterium adolescentis L2-32 | GCA_000154085.1 |
| Bifidobacterium bifidum ATCC 29521 | AP012323.1 |
| Bifidobacterium pseudocatenulatum DSM 20438 | GCA_000173435.1 |
| Blautia hansenii DSM 20583 | CP022413.2 |
| Blautia luti DSM 14534 | GCA_009707925.1 |
| Candidatus Ruthia magnifica str. | CP000488.1 |
| Cereibacter sphaeroides ATCC 17029 | GCA_000015985.1 |
| Citrobacter youngae ATCC 29220 | GCA_000155975.1 |
| Clostridium bolteae ATCC BAA-613 | CP022464.2/CP022465.2 |
| Clostridium sp. M62/1 | GCA_000159055.1 |
| Clostridium sporogenes ATCC 15579 | GCA_000155085.1 |
| Collinsella intestinalis DSM 13280 | GCA_000156175.1 |
| Collinsella stercoris DSM 13279 | GCA_000156215.1 |
| Coprococcus comes ATCC 27758 | GCA_000155875.1 |
| Dorea formicigenerans ATCC 27755 | GCA_000169235.1 |
| Edwardsiella tarda ATCC 23685 | GCA_000163955.1 |
| Enterobacter cancerogenus ATCC 35316 | GCA_000155995.1 |
| Escherichia fergusonii ATCC 35469 | GCA_000026225.1 |
| Geotalea uraniireducens Rf4 | CP000698.1 |
| Gluconacetobacter diazotrophicus PA1 5 | GCA_000021325.1 |
| Granulicella tundricola MP5ACTX9 | GCA_000178975.2 |
| Herpetosiphon aurantiacus DSM 785 | GCA_000018565.1 |
| Holdemanella biformis DSM 3989 | GCA_000156655.1 |
| Holdemania filiformis DSM 12042 | GCA_000157995.1 |
| Hungatella hathewayi DSM 13479 | GCA_000160095.1 |
| Lachnospira eligens ATCC 27750 | GCA_000146185.1 |
| Lacticaseibacillus paracasei ATCC 334 | GCA_000014525.1 |
| Lactobacillus reuteri DSM 20016 | CP000705.1 |
| Lactococcus lactis subsp. Lactis | GCA_003176835.1 |
| Leptothrix cholodnii SP-6 | CP001013.1 |
| Ligilactobacillus ruminis DSM 20403 = NBRC 102161 | GCA_001436475.1 |
| Maricaulis maris MCS10 | CP000449.1 |
| Marvinbryantia formatexigens DSM 14469 | GCA_000173815.1 |
| Megamonas funiformis YIT 11815 | GCA_000245775.1 |
| Methylorubrum extorquens CM4 | GCA_000021845.1 |
| Methylovorus glucosotrophus | GCA_000023745.1 |
| Nitratifractor salsuginis DSM 16511 | CP002452.1 |
| Nocardiopsis dassonvillei subsp. dassonvillei DSM 43111 | GCA_000092985.1 |
| Parabacteroides johnsonii DSM 18315 | GCA_000156495.1 |
| Parabacteroides merdae ATCC 43184 | GCA_000154105.1 |
| Pelodictyon phaeoclathratiforme BU-1 | CP001110.1 |
| Phocaeicola dorei DSM 17855 | GCA_000156075.1 |
| Polynucleobacter necessarius subsp. necessarius STIR1 | CP001010.1 |
| Porphyromonas asaccharolytica DSM 20707 | CP002689.1 |
| Proteus penneri ATCC 35198 | GCA_000155835.1 |
| Rhizobium leguminosarum bv. trifolii WSM1325 | GCA_000023185.1 |
| Roseburia intestinalis L1-82 | GCA_000156535.1 |
| Ruminiclostridium cellulolyticum H10 | CP001348.1 |
| Ruminococcus gnavus ATCC 29149 | GCA_000169475.1 |
| Salinispora arenicola CNS-205 | CP000850.1 |
| Sediminispirochaeta smaragdinae DSM 11293 | CP002116.1 |
| Shewanella baltica OS117 | GCA_000215895.1 |
| Solitalea canadensis DSM 3403 | CP003349.1 |
| Sphaerobacter thermophilus DSM 20745 | GCA_000024985.1 |
| Sphaerochaeta pleomorpha str. Grapes | CP003155.1 |
| Stanieria cyanosphaera PCC 7437 | GCA_000317575.1 |
| Staphylococcus aureus subsp. aureus JH9 | GCA_000016805.1 |
| Streptococcus infantarius ATCC BAA-102 | GCA_000154985.1 |
| Subdoligranulum variabile DSM 15176 | GCA_000157955.1 |
| Syntrophobacter fumaroxidans MPOB | CP000478.1 |
| Thermoanaerobacterium thermosaccharolyticum DSM 571 | CP002171.1 |
| Thermus thermophilus SG0.5JP17-16 | GCA_000214845.1 |
| Thomasclavelia ramosa DSM 1402 | GCA_000154485.1 |
| Trichormus variabilis ATCC 29413 | GCA_000204075.1 |
| Tsukamurella paurometabola DSM 20162 | GCA_000092225.1 |
| Weeksella virosa DSM 16922 | CP002455.1 |
| Xylanimonas cellulosilytica DSM 15894 | GCA_000024965.1 |

**Table S22** List of organisms in Sim92 dataset

| **Organisms** | **Accession** |
| --- | --- |
| [Clostridium] asparagiforme DSM 15981 | GCA_000158075.1 |
| [Clostridium] hylemonae DSM 15053 | GCA_000156515.1 |
| [Clostridium] nexile DSM 1787 | GCA_000156035.2 |
| [Clostridium] symbiosum ATCC 14940 | GCA_000466485.1 |
| Acetivibrio thermocellus DSM 2360 | CP016502.1 |
| Acidobacterium capsulatum ATCC 51196 | CP001472.1 |
| Alistipes indistinctus YIT 12060 | JH370372.1 |
| Bacteroides cellulosilyticus DSM 14838 | GCA_000158035.1 |
| Bacteroides finegoldii DSM 17565 | GCA_000156195.1 |
| Bacteroides intestinalis DSM 17393 | GCA_000172175.1 |
| Bacteroides ovatus strain ATCC 8483 | CP012938.1 |
| Bacteroides plebeius DSM 17135 | GCA_000187895.1 |
| Bacteroides stercoris ATCC 43183 | GCA_000154525.1 |
| Bacteroides thetaiotaomicron 3731 | GCA_014131755.1 |
| Bacteroides thetaiotaomicron 7330 | CP012937.1 |
| Bacteroides thetaiotaomicron strain DSM 2079 | CP040530.1 |
| Bacteroides thetaiotaomicron VPI-5482 | GCA_000011065.1 |
| Bacteroides uniformis ATCC 8492 | GCA_000154205.1 |
| Bacteroides vulgatus ATCC 8482 | CP000139.1 |
| Bifidobacterium adolescentis L2-32 | GCA_000154085.1 |
| Bifidobacterium angulatum DSM 20098 = JCM 7096 | AP012322.1 |
| Bifidobacterium bifidum ATCC 29521 | AP012323.1 |
| Bifidobacterium pseudocatenulatum DSM 20438 | GCA_000173435.1 |
| Blautia hansenii DSM 20583 | CP022413.2 |
| Blautia luti DSM 14534 | GCA_009707925.1 |
| Caldicellulosiruptor bescii DSM 6725 | GCA_000022325.1 |
| Caldicellulosiruptor saccharolyticus DSM 8903 | CP000679.1 |
| Candidatus Aciduliprofundum boonei | GCA_013329755.1 |
| Chlorobium phaeovibrioides | GCA_003968655.1 |
| Citrobacter youngae ATCC 29220 | GCA_000155975.1 |
| Clostridium bolteae ATCC BAA-613 | CP022464.2/CP022465.2 |
| Clostridium sp. M62/1 | GCA_000159055.1 |
| Collinsella intestinalis DSM 13280 | GCA_000156175.1 |
| Collinsella stercoris DSM 13279 | GCA_000156215.1 |
| Coprococcus comes ATCC 27758 | GCA_000155875.1 |
| Deinococcus radiodurans R1 = ATCC 13939 = DSM 20539 strain | CP068791.1 |
| Desulfovibrio piger | GCA_937907355.1 |
| Desulfovibrio vulgaris str. 'Miyazaki F' | CP001197.1 |
| Dictyoglomus turgidum DSM 6724 | CP001251.1 |
| Dorea formicigenerans ATCC 27755 | GCA_000169235.1 |
| Edwardsiella tarda ATCC 23685 | GCA_000163955.1 |
| Enterobacter cancerogenus ATCC 35316 | GCA_000155995.1 |
| Enterococcus faecalis EnGen0336 | GCA_000393015.1 |
| Escherichia fergusonii ATCC 35469 | GCA_000026225.1 |
| Haloferax volcanii DS2 | GCA_000337315.1 |
| Herpetosiphon aurantiacus DSM 785 | GCA_000018565.1 |
| Holdemanella biformis DSM 3989 | GCA_000156655.1 |
| Holdemania filiformis DSM 12042 | GCA_000157995.1 |
| Hungatella hathewayi DSM 13479 | GCA_000160095.1 |
| Hydrogenobaculum sp. Y04AAS1 | NC_011126.1 |
| Lachnospira eligens ATCC 27750 | GCA_000146185.1 |
| Ligilactobacillus ruminis DSM 20403 = NBRC 102161 | GCA_001436475.1 |
| Marvinbryantia formatexigens DSM 14469 | GCA_000173815.1 |
| Megamonas funiformis YIT 11815 | GCA_000245775.1 |
| Methanocaldococcus jannaschii DSM 2661 | GCA_000091665.1 |
| Methanococcus maripaludis C5 | GCA_000016125.1 |
| Methanococcus maripaludis strain S2 | BX950229.1 |
| Methanopyrus kandleri AV19 | AE009439.1 |
| Nanoarchaeum equitans Kin4-M | AE017199.1 |
| Nitrosomonas europaea ATCC 19718 | AL954747.1 |
| Nostoc sp. PCC 7120 = FACHB-418 | GCA_003990585.1 |
| Parabacteroides johnsonii DSM 18315 | GCA_000156495.1 |
| Parabacteroides merdae ATCC 43184 | GCA_000154105.1 |
| Paraburkholderia xenovorans LB400 | GCA_000756045.1 |
| Persephonella marina EX-H1 | GCA_000021565.1 |
| Phocaeicola dorei DSM 17855 | GCA_000156075.1 |
| Phocaeicola vulgatus MG01-10 DNA | AP025240.1 |
| Porphyromonas gingivalis ATCC 33277 | AP009380.1 |
| Proteus penneri ATCC 35198 | GCA_000155835.1 |
| Pyrobaculum aerophilum str. IM2 | AE009441.1 |
| Pyrobaculum arsenaticum DSM 13514 | CP000660.1 |
| Pyrobaculum calidifontis JCM 11548 | CP000561.1 |
| Pyrococcus furiosus DSM 3638 | CP023154.1 |
| Roseburia intestinalis L1-82 | GCA_000156535.1 |
| Ruegeria pomeroyi DSS-3 | CP000031.2 |
| Ruminococcus gnavus ATCC 29149 | GCA_000169475.1 |
| Shewanella baltica OS185 | GCA_000017325.1 |
| Shewanella baltica OS223 | GCA_000021665.1 |
| Streptococcus infantarius ATCC BAA-102 | GCA_000154985.1 |
| Subdoligranulum variabile DSM 15176 | GCA_000157955.1 |
| Sulfitobacter sp. EE-36 | GCA_000152605.1 |
| Sulfitobacter sp. NAS-14.1 | GCA_000152645.1 |
| Sulfolobus tokodaii str. 7 DNA | BA000023.2 |
| Sulfurihydrogenibium yellowstonense SS-5 | GCA_000173615.1 |
| Thermoanaerobacter pseudethanolicus ATCC 33223 | CP000924.1 |
| Thermotoga neapolitana DSM 4359 | CP000916.1 |
| Thermotoga petrophila RKU-1 | CP000702.1 |
| Thermotoga sp. RQ2 | NC_010483.1 |
| Thermus thermophilus HB8 | GCA_000091545.1 |
| Thomasclavelia ramosa DSM 1402 | GCA_000154485.1 |
| Treponema denticola ATCC 35405 | AE017226.1 |
| Wolinella succinogenes DSM 1740 | BX571656.1 |

**Table S23** List of organisms in Sim98 dataset

| **Organisms** | **Accession** |
| --- | --- |
| [Clostridium] asparagiforme DSM 15981 | GCA_000158075.1 |
| [Clostridium] hylemonae DSM 15053 | GCA_000156515.1 |
| [Clostridium] nexile DSM 1787 | GCA_000156035.2 |
| [Clostridium] symbiosum ATCC 14940 | GCA_000466485.1 |
| Akkermansia muciniphila | GCA_009731575.1 |
| Akkermansia muciniphila ATCC BAA-835 | GCA_017504145.1 |
| Alistipes finegoldii DSM 17242 | NC_018011.1 |
| Alistipes indistinctus YIT 12060 | JH370372.1 |
| Anaerobutyricum hallii DSM 3353 | GCA_000173975.1 |
| Anaerostipes hadrus ATCC 29173 = JCM 17467 | NZ_KB290627.1 |
| Bacteroides cellulosilyticus DSM 14838 | GCA_000158035.1 |
| Bacteroides finegoldii DSM 17565 | GCA_000156195.1 |
| Bacteroides intestinalis DSM 17393 | GCA_000172175.1 |
| Bacteroides ovatus strain ATCC 8483 | CP012938.1 |
| Bacteroides plebeius DSM 17135 | GCA_000187895.1 |
| Bacteroides stercoris ATCC 43183 | GCA_000154525.1 |
| Bacteroides thetaiotaomicron 3731 | GCA_014131755.1 |
| Bacteroides thetaiotaomicron 7330 | CP012937.1 |
| Bacteroides thetaiotaomicron strain DSM 2079 | CP040530.1 |
| Bacteroides thetaiotaomicron VPI-5482 | GCA_000011065.1 |
| Bacteroides uniformis ATCC 8492 | GCA_000154205.1 |
| Bacteroides vulgatus ATCC 8482 | CP000139.1 |
| Bifidobacterium adolescentis L2-32 | GCA_000154085.1 |
| Bifidobacterium longum subsp. longum JCM 1217 | GCA_000196555.1 |
| Bifidobacterium pseudocatenulatum DSM 20438 | GCA_000173435.1 |
| Blautia hansenii DSM 20583 | CP022413.2 |
| Blautia luti DSM 14534 | GCA_009707925.1 |
| Blautia wexlerae DSM 19850 | GCA_025148125.1 |
| Caldicellulosiruptor bescii DSM 6725 | GCA_000022325.1 |
| Caldicellulosiruptor saccharolyticus DSM 8903 | CP000679.1 |
| Candidatus Aciduliprofundum boonei | GCA_013329755.1 |
| Chlorobium phaeobacteroides DSM 266 | GCA_000015125.1 |
| Chlorobium phaeovibrioides | GCA_003968655.1 |
| Chlorobium tepidum TLS | AE006470.1 |
| Chloroflexus aurantiacus J-10-fl | GCA_000018865.1 |
| Citrobacter youngae ATCC 29220 | GCA_000155975.1 |
| Clostridium bolteae ATCC BAA-613 | CP022464.2/CP022465.2 |
| Clostridium butyricum DSM 10702 | GCA_000409755.1 |
| Clostridium sp. M62/1 | GCA_000159055.1 |
| Collinsella aerofaciens | GCA_002736145.1 |
| Collinsella intestinalis DSM 13280 | GCA_000156175.1 |
| Collinsella stercoris DSM 13279 | GCA_000156215.1 |
| Coprococcus comes ATCC 27758 | GCA_000155875.1 |
| Deinococcus radiodurans R1 = ATCC 13939 = DSM 20539 strain | CP068791.1 |
| Desulfovibrio piger | GCA_937907355.1 |
| Dictyoglomus turgidum DSM 6724 | CP001251.1 |
| Dorea formicigenerans ATCC 27755 | GCA_000169235.1 |
| Edwardsiella tarda ATCC 23685 | GCA_000163955.1 |
| Enterobacter cancerogenus ATCC 35316 | GCA_000155995.1 |
| Enterococcus faecalis EnGen0336 | GCA_000393015.1 |
| Escherichia fergusonii ATCC 35469 | GCA_000026225.1 |
| Fusobacterium nucleatum subsp. | LN831027.1 |
| Haloferax volcanii DS2 | GCA_000337315.1 |
| Herpetosiphon aurantiacus DSM 785 | GCA_000018565.1 |
| Holdemanella biformis DSM 3989 | GCA_000156655.1 |
| Holdemania filiformis DSM 12042 | GCA_000157995.1 |
| Hungatella hathewayi DSM 13479 | GCA_000160095.1 |
| Hydrogenobaculum sp. Y04AAS1 | NC_011126.1 |
| Lachnospira eligens ATCC 27750 | GCA_000146185.1 |
| Lactobacillus gasseri ATCC 33323 = JCM 1131 | NC_008530.1 |
| Lactobacillus reuteri DSM 20016 | CP000705.1 |
| Ligilactobacillus ruminis DSM 20403 = NBRC 102161 | GCA_001436475.1 |
| Marvinbryantia formatexigens DSM 14469 | GCA_000173815.1 |
| Megamonas funiformis YIT 11815 | GCA_000245775.1 |
| Methanocaldococcus jannaschii DSM 2661 | GCA_000091665.1 |
| Methanococcus maripaludis strain S2 | BX950229.1 |
| Nanoarchaeum equitans Kin4-M | AE017199.1 |
| Nitrosomonas europaea ATCC 19718 | AL954747.1 |
| Nostoc sp. PCC 7120 = FACHB-418 | GCA_003990585.1 |
| Parabacteroides johnsonii DSM 18315 | GCA_000156495.1 |
| Parabacteroides merdae ATCC 43184 | GCA_000154105.1 |
| Paraburkholderia xenovorans LB400 | GCA_000756045.1 |
| Phocaeicola dorei DSM 17855 | GCA_000156075.1 |
| Phocaeicola vulgatus MG01-10 DNA | AP025240.1 |
| Prevotella melaninogenica ATCC 25845 | CP002123.1 |
| Proteus penneri ATCC 35198 | GCA_000155835.1 |
| Pyrococcus horikoshii OT3 | BA000001.2 |
| Roseburia intestinalis L1-82 | GCA_000156535.1 |
| Ruegeria pomeroyi DSS-3 | CP000031.2 |
| Ruminococcus gauvreauii DSM 19829 | GCA_000425525.1 |
| Ruminococcus gnavus ATCC 29149 | GCA_000169475.1 |
| Salinispora tropica CNB-440 | GCA_000016425.1 |
| Segatella copri DSM 18205 | GCA_020735445.1 |
| Shewanella baltica OS185 | GCA_000017325.1 |
| Shewanella baltica OS223 | GCA_000021665.1 |
| Streptococcus infantarius ATCC BAA-102 | GCA_000154985.1 |
| Subdoligranulum variabile DSM 15176 | GCA_000157955.1 |
| Sulfitobacter sp. EE-36 | GCA_000152605.1 |
| Sulfitobacter sp. NAS-14.1 | GCA_000152645.1 |
| Sulfurihydrogenibium sp. YO3AOP1 | NC_010730.1 |
| Sulfurihydrogenibium yellowstonense SS-5 | GCA_000173615.1 |
| Thermoanaerobacter pseudethanolicus ATCC 33223 | CP000924.1 |
| Thermotoga neapolitana DSM 4359 | CP000916.1 |
| Thermotoga petrophila RKU-1 | CP000702.1 |
| Thermotoga sp. RQ2 | NC_010483.1 |
| Thermus thermophilus HB8 | GCA_000091545.1 |
| Thomasclavelia ramosa DSM 1402 | GCA_000154485.1 |
| Wolinella succinogenes DSM 1740 | BX571656.1 |

**Table S24** AMBER results for simulated datasets

| **Dataset** | **Ground**  **truth**  **bins** | **Tool** | **Number of bins identified** | **Accuracy (seq)** | **Purity (seq)** | **Completeness (seq)** | **F1 score for sample (seq)** | **Rand Index (seq)** |
| --- | --- | --- | --- | --- | --- | --- | --- | --- |
| Sim-5G | 5 | BusyBee Web | 2 | 0.431 | 0.431 | **0.956** | 0.595 | 0.586 |
|  |  | CONCOCT | 9 | **0.923** | 0.923 | 0.790 | 0.852 | 0.889 |
|  |  | MaxBin 2.0 | **5** | 0.915 | 0.915 | 0.915 | **0.915** | 0.919 |
|  |  | MetaBAT 2 | **5** | 0.665 | **1.00** | 0.665 | 0.799 | **1.00** |
|  |  | MetaDecoder | **5** | 0.839 | 0.972 | 0.839 | 0.900 | 0.979 |
|  |  | CoCoBin | 10 | 0.839 | 0.986 | 0.569 | 0.721 | 0.879 |
| SRR8304764 | 51 | BusyBee Web | 29 | 0.489 | 0.489 | **0.799** | 0.607 | 0.911 |
|  |  | CONCOCT | 77 | 0.521 | 0.521 | 0.604 | 0.559 | 0.954 |
|  |  | MaxBin 2.0 | 35 | 0.448 | 0.577 | 0.512 | 0.542 | 0.957 |
|  |  | MetaBAT 2 | 64 | 0.333 | **0.733** | 0.378 | 0.499 | 0.970 |
|  |  | MetaDecoder | 77 | 0.415 | 0.680 | 0.351 | 0.463 | **0.975** |
|  |  | CoCoBin | **51** | **0.542** | 0.616 | 0.685 | **0.649** | 0.965 |
| SRR8304773 | 51 | BusyBee Web | 20 | 0.386 | 0.386 | **0.837** | 0.529 | 0.839 |
|  |  | CONCOCT | 78 | **0.562** | 0.563 | 0.604 | 0.583 | 0.956 |
|  |  | MaxBin 2.0 | 41 | 0.519 | 0.601 | 0.545 | 0.572 | 0.955 |
|  |  | MetaBAT 2 | 88 | 0.303 | **0.788** | 0.295 | 0.429 | 0.970 |
|  |  | MetaDecoder | 71 | 0.399 | 0.741 | 0.376 | 0.499 | **0.976** |
|  |  | CoCoBin | **55** | 0.536 | 0.638 | 0.603 | **0.620** | 0.965 |
| SRR8304775 | 51 | BusyBee Web | 20 | 0.477 | 0.477 | **0.764** | 0.588 | 0.908 |
|  |  | CONCOCT | 84 | 0.615 | 0.616 | 0.658 | 0.636 | 0.950 |
|  |  | MaxBin 2.0 | 32 | 0.527 | 0.538 | 0.647 | 0.588 | 0.947 |
|  |  | MetaBAT 2 | 42 | 0.234 | 0.903 | 0.233 | 0.370 | 0.983 |
|  |  | MetaDecoder | 30 | 0.290 | **0.919** | 0.289 | 0.439 | **0.988** |
|  |  | CoCoBin | **47** | **0.620** | 0.726 | 0.690 | **0.708** | 0.965 |
| SRR8304776 | 51 | BusyBee Web | 16 | 0.391 | 0.391 | **0.811** | 0.528 | 0.898 |
|  |  | CONCOCT | 80 | **0.593** | 0.594 | 0.663 | 0.627 | 0.941 |
|  |  | MaxBin 2.0 | 31 | 0.529 | 0.548 | 0.643 | 0.592 | 0.945 |
|  |  | MetaBAT 2 | 46 | 0.244 | **0.900** | 0.232 | 0.369 | **0.983** |
|  |  | MetaDecoder | 36 | 0.308 | 0.882 | 0.290 | 0.436 | **0.983** |
|  |  | CoCoBin | **48** | 0.585 | 0.688 | 0.706 | **0.697** | 0.964 |
| Sim79 | 79 | BusyBee Web | 31 | 0.517 | 0.517 | **0.806** | **0.630** | 0.931 |
|  |  | CONCOCT | 75 | **0.547** | 0.548 | 0.575 | 0.561 | 0.956 |
|  |  | MaxBin 2.0 | 49 | 0.327 | 0.345 | 0.445 | 0.388 | 0.946 |
|  |  | MetaBAT 2 | 109 | 0.172 | **0.829** | 0.137 | 0.236 | 0.975 |
|  |  | MetaDecoder | 52 | 0.252 | 0.781 | 0.243 | 0.370 | **0.980** |
|  |  | CoCoBin | **82** | 0.494 | 0.640 | 0.544 | 0.588 | 0.969 |
| Sim82 | 82 | BusyBee Web | 40 | **0.615** | 0.615 | **0.822** | **0.704** | 0.959 |
|  |  | CONCOCT | 76 | 0.580 | 0.580 | 0.601 | 0.591 | 0.961 |
|  |  | MaxBin 2.0 | 45 | 0.396 | 0.406 | 0.538 | 0.463 | 0.948 |
|  |  | MetaBAT 2 | 118 | 0.148 | **0.800** | 0.127 | 0.219 | **0.979** |
|  |  | MetaDecoder | 55 | 0.217 | 0.769 | 0.214 | 0.335 | **0.979** |
|  |  | CoCoBin | **86** | 0.550 | 0.673 | 0.633 | 0.652 | 0.974 |
| Sim92 | 92 | BusyBee Web | 30 | 0.464 | 0.464 | **0.827** | **0.594** | 0.913 |
|  |  | CONCOCT | 86 | **0.538** | 0.539 | 0.577 | 0.557 | 0.953 |
|  |  | MaxBin 2.0 | 43 | 0.289 | 0.296 | 0.462 | 0.361 | 0.940 |
|  |  | MetaBAT 2 | 123 | 0.149 | **0.854** | 0.100 | 0.179 | **0.980** |
|  |  | MetaDecoder | 59 | 0.255 | 0.743 | 0.241 | 0.363 | 0.977 |
|  |  | CoCoBin | **96** | 0.504 | 0.653 | 0.533 | 0.587 | 0.966 |
| Sim98 | 98 | BusyBee Web | 30 | 0.396 | 0.396 | **0.782** | **0.525** | 0.889 |
|  |  | CONCOCT | **90** | **0.471** | 0.472 | 0.533 | 0.500 | 0.946 |
|  |  | MaxBin 2.0 | 45 | 0.236 | 0.248 | 0.389 | 0.303 | 0.948 |
|  |  | MetaBAT 2 | 122 | 0.092 | **0.815** | 0.056 | 0.105 | **0.976** |
|  |  | MetaDecoder | 65 | 0.234 | 0.664 | 0.222 | 0.333 | 0.972 |
|  |  | CoCoBin | 107 | 0.399 | 0.531 | 0.453 | 0.489 | 0.965 |

The best result in each category is red bold. The second-best result in each category is underlined.

**Table S25** AMBER results for mock community datasets.

| Dataset | Ground  truth  bins | Tool | Number of bins identified | Accuracy (seq) | Purity (seq) | Completeness (seq) | F1 score for sample (seq) | Rand Index (seq) |
| --- | --- | --- | --- | --- | --- | --- | --- | --- |
| SRR8073716 | 12 | BusyBee Web | 5 | 0.354 | 0.424 | **0.719** | 0.533 | 0.840 |
|  |  | CONCOCT | 16 | **0.557** | 0.666 | 0.605 | **0.634** | 0.891 |
|  |  | MaxBin 2.0 | **10** | 0.482 | 0.578 | 0.592 | 0.585 | 0.876 |
|  |  | MetaBAT 2 | 20 | 0.325 | **0.978** | 0.264 | 0.416 | **0.960** |
|  |  | MetaDecoder | **14** | 0.466 | 0.769 | 0.468 | 0.581 | 0.908 |
|  |  | CoCoBin | 20 | 0.525 | 0.730 | 0.423 | 0.536 | 0.906 |
| SRR11487931 | 20 | BusyBee Web | 12 | 0.819 | 0.819 | 0.842 | 0.831 | 0.934 |
|  |  | CONCOCT | 37 | **0.882** | 0.882 | **0.876** | **0.879** | 0.962 |
|  |  | MaxBin 2.0 | 11 | 0.686 | 0.693 | 0.840 | 0.759 | 0.902 |
|  |  | MetaBAT 2 | 12 | 0.274 | **0.980** | 0.273 | 0.427 | **0.990** |
|  |  | MetaDecoder | 9 | 0.234 | 0.932 | 0.223 | 0.359 | 0.960 |
|  |  | CoCoBin | **22** | 0.844 | 0.946 | 0.757 | 0.841 | 0.962 |
| SRR11487935 | 20 | BusyBee Web | 11 | 0.807 | 0.807 | 0.839 | 0.823 | 0.936 |
|  |  | CONCOCT | 28 | **0.881** | 0.882 | **0.886** | **0.884** | 0.968 |
|  |  | MaxBin 2.0 | 11 | 0.703 | 0.709 | 0.835 | 0.767 | 0.903 |
|  |  | MetaBAT 2 | 12 | 0.227 | **0.969** | 0.224 | 0.364 | **0.978** |
|  |  | MetaDecoder | 11 | 0.225 | 0.886 | 0.229 | 0.364 | 0.944 |
|  |  | CoCoBin | **18** | 0.837 | 0.941 | 0.785 | 0.856 | 0.967 |
| SRR3656745 | 26 | BusyBee Web | 8 | 0.416 | 0.416 | **0.909** | 0.570 | 0.894 |
|  |  | CONCOCT | 30 | 0.584 | 0.584 | 0.803 | 0.676 | 0.927 |
|  |  | MaxBin 2.0 | **26** | 0.660 | 0.663 | 0.829 | 0.737 | 0.951 |
|  |  | MetaBAT 2 | 38 | 0.660 | **0.965** | 0.604 | 0.743 | **0.977** |
|  |  | MetaDecoder | **26** | 0.702 | 0.914 | 0.700 | 0.793 | 0.974 |
|  |  | CoCoBin | 24 | **0.750** | 0.881 | 0.775 | **0.825** | 0.969 |
| SRR606249 | 64 | BusyBee Web | 28 | 0.690 | 0.690 | **0.947** | 0.799 | 0.971 |
|  |  | CONCOCT | 71 | **0.819** | 0.819 | 0.872 | **0.845** | 0.980 |
|  |  | MaxBin 2.0 | 55 | 0.748 | 0.757 | 0.761 | 0.759 | 0.974 |
|  |  | MetaBAT 2 | 74 | 0.507 | **0.933** | 0.514 | 0.663 | **0.985** |
|  |  | MetaDecoder | 71 | 0.551 | 0.910 | 0.511 | 0.655 | **0.984** |
|  |  | CoCoBin | **59** | 0.788 | 0.839 | 0.849 | 0.844 | 0.976 |

The best result in each category is red bold. The second-best result in each category is underlined.

Table S26 AMBER results for real-world (Sharon) dataset

| Dataset | Number of bins | Tool | Number of bins identified | Accuracy (seq) | Purity (seq) | Completeness (seq) | F1 score for sample (seq) | Rand Index (seq) |
| --- | --- | --- | --- | --- | --- | --- | --- | --- |
| **Sharon** | 32 | BusyBee Web | 6 | 0.540 | 0.540 | **0.952** | 0.689 | 0.733 |
|  |  | CONCOCT | 45 | **0.823** | 0.824 | 0.719 | **0.768** | 0.880 |
|  |  | MaxBin 2.0 | 12 | 0.510 | 0.510 | 0.641 | 0.568 | 0.699 |
|  |  | MetaBAT 2 | 12 | 0.377 | **0.942** | 0.391 | 0.553 | **0.963** |
|  |  | MetaDecoder | 9 | 0.413 | 0.899 | 0.246 | 0.386 | 0.676 |
|  |  | CoCoBin | **29** | 0.799 | 0.870 | 0.572 | 0.690 | 0.850 |

The best result in each category is red bold. The second-best result in each category is underlined.
